# Repeated fasting events sensitize enhancers, transcription factor activity and gene expression to support augmented ketogenesis

**DOI:** 10.1101/2024.05.07.592891

**Authors:** Noga Korenfeld, Meital Charni-Natan, Justine Bruse, Dana Goldberg, Dorin Marciano-Anaki, Dan Rotaro, Tali Gorbonos, Talia Radushkevitz-Frishman, Arnaud Polizzi, Abed Nasereddin, Meirav Bar-Shimon, Anne Fougerat, Hervé Guillou, Ido Goldstein

**Affiliations:** Institute of Biochemistry, Food Science and Nutrition. The Robert H. Smith Faculty of Agriculture, Food and Environment. The Hebrew University of Jerusalem. POB 12, Rehovot 7610001, Israel; Toxalim (Research Center in Food Toxicology), INRAE, ENVT, INP-PURPAN, UMR 1331, UPS, Université de Toulouse, Toulouse, France; Genomics Applications Laboratory, Core Research Facility, Faculty of Medicine, The Hebrew University of Jerusalem-Hadassah Medical School

**Keywords:** Fasting, Enhancers, Transcriptional regulation, Transcription factors, Nuclear Receptors, Metabolism, Chromatin

## Abstract

Mammals withstand frequent and prolonged fasting periods due to hepatic production of ketone bodies. Because the fasting response is transcriptionally-regulated, we asked whether enhancer dynamics impose a transcriptional program during recurrent fasting and whether this generates effects distinct from a single fasting bout. We found that mice undergoing alternate-day fasting (ADF) respond profoundly differently to a following fasting bout compared to mice first experiencing fasting. Hundreds of genes enabling ketogenesis are ‘sensitized’ (induced more strongly by fasting following ADF). Liver enhancers regulating these genes are also sensitized and harbor increased binding of PPARα, the main ketogenic transcription factor. ADF leads to augmented ketogenesis compared to a single fasting bout in wild-type, but not hepatocyte-specific PPARα-deficient mice. Thus, we found that past fasting events are ‘remembered’ in hepatocytes, sensitizing their enhancers to the next fasting bout and augment ketogenesis. Our findings shed light on transcriptional regulation mediating adaptation to repeated signals.

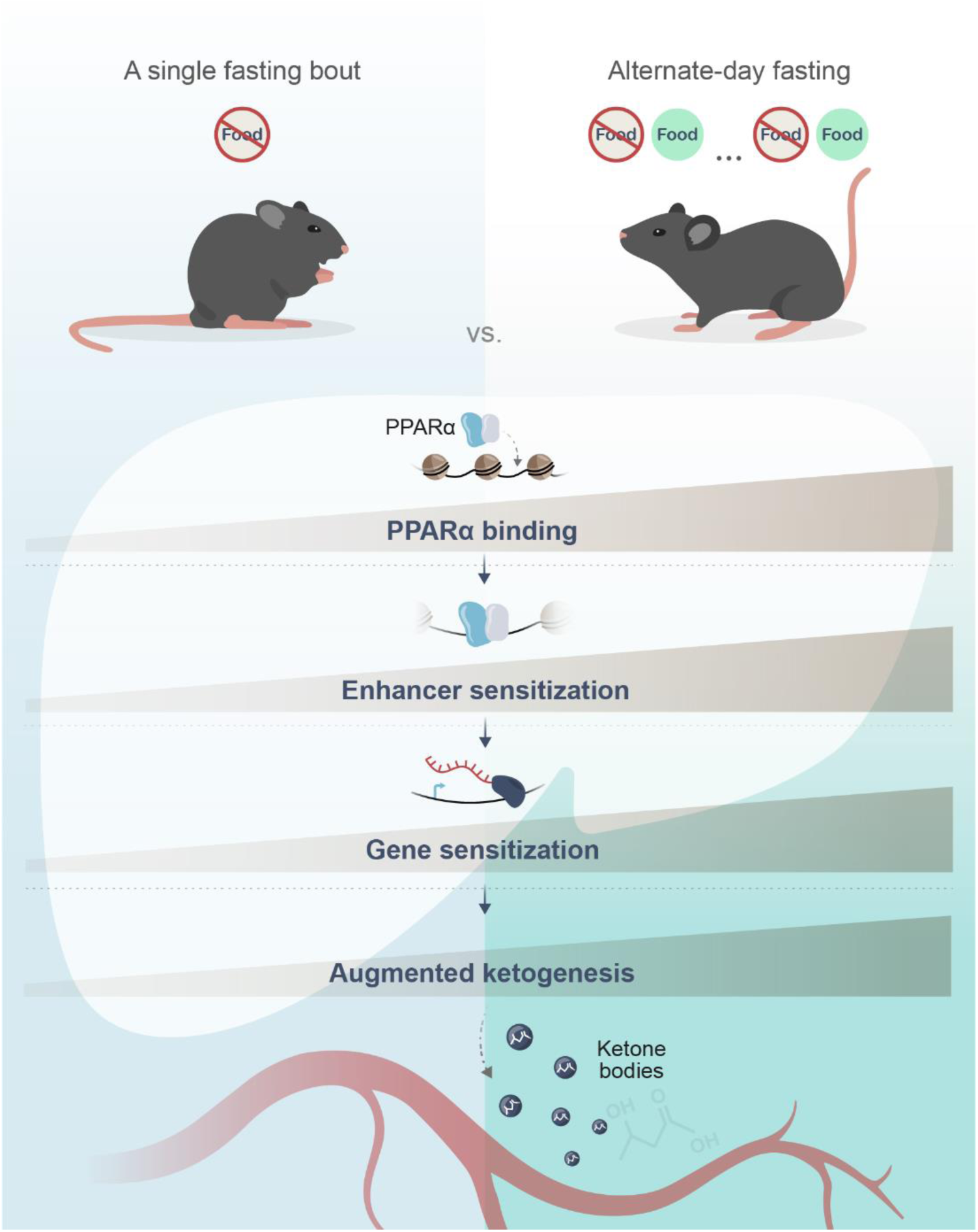

## Introduction

Episodes of fasting are an inherent aspect of physiology with most animals experiencing frequent and sometimes prolonged bouts of fasting^1^. Mammals are exquisitely fitted to tolerate extended periods without food intake due to an adaptive response to fasting. The fasting response includes a diverse set of endocrine, metabolic and neural cues that together affect metabolism and behavior to maintain homeostasis in the face of energy shortage. A central aspect of the fasting response is the hepatic production of fuels in the form of glucose and ketone bodies that supply the energetic needs of extra-hepatic tissues. Glucose is produced by both glycogenolysis (the breaking down of glycogen) and gluconeogenesis, the de novo synthesis of glucose from precursors such as amino acids, lactate and glycerol. In ketogenesis, acetyl-CoA is used as a precursor to producing ketone bodies, mainly beta-hydroxybutyrate (BHB). An abundant supply of acetyl-CoA is achieved by lipolysis of triglycerides in adipose tissue, the release of free fatty acids to circulation, their uptake by hepatocytes and fatty acid oxidation (FAO) into acetyl-CoA. Glycogen breakdown is a quick and efficient way to supply fuel but hepatic glycogen depots quickly dwindle and are thus not a principal source of fuel during prolonged fasting bouts. In contrast, both gluconeogenesis and especially ketogenesis serve to provide fuel during prolonged fasting^2–5^.

Hepatic fuel production is heavily regulated at the transcriptional levels. Hundreds of genes related to gluconeogenesis, FAO and ketogenesis as well as to cellular processes enabling these pathways (uptake of extracellular precursors, inter-organelle transport, lipid catabolism etc.) are transcriptionally regulated during fasting. This widespread regulation of gene transcription is mediated by various transcription factors (TFs), some of which are activated by fasting-related extracellular signals such as hormones and metabolites. These TFs include cAMP response element binding protein (CREB) which is activated by glucagon signaling, peroxisome proliferator-activated receptor alpha (PPARα) which is activated by fatty acids, glucocorticoid receptor (GR) which is activated by glucocorticoids and forkhead box O1 (FoxO1) which is activated by decreased insulin levels during fasting^6–8^. TFs bind cis-regulatory DNA elements termed enhancers. Upon activation and enhancer binding, TFs recruit histone-modifying enzymes, co-activators and chromatin-remodeling enzymes that together increase enhancer accessibility and promote gene transcription. Therefore, an increase in accessibility is often used as a proxy for increased enhancer activity^9,10^. In addition to causing a bulk increase in enhancer accessibility, TFs also leave a ‘footprint’ on their binding site within the enhancer (measured by local protection from nuclease cleavage)^11^. During fasting, thousands of enhancers are activated by fasting-related TFs, mediating the transcriptional response to fasting^12^. These activated enhancers show abundant TF binding, TF footprints and increased accessibility, all resulting in fasting-dependent gene induction^6,12,13^.

A multitude of studies in mammals, especially rodents and humans, examined the effects of nutritional regimens that involve recurring fasting events (i.e., intermittent fasting). The most extensively-studied nutritional regimens are alternate-day fasting (ADF) and time-restricted feeding. In ADF, food is consumed ad libitum for 24 h followed by a complete lack of food consumption (or severely restricted food consumption) for the next 24 h. In time-restricted feeding, within a 24 h window a fasting period of around 18 h is imposed following a shorter period (around 6 h) of ad libitum feeding. In each of these regimens, far-reaching health benefits were reported in rodents and humans. These benefits include improvements in glucose tolerance, insulin sensitivity and lipid profiles as well as weight loss. Also, amelioration of obesity, diabetes, steatosis, hypertension, inflammation, certain cancers and neurodegenerative diseases were observed (reviewed in^14–17^). A concern was raised that intermittent fasting regimens lead to an overall reduction in calorie intake, raising the possibility that the observed health benefits stem from the ensuing caloric restriction rather than from fasting per se. However, data has accumulated to show that the metabolic/hormonal/physiological state that fasting imposes leads to the striking health benefits rather than mere caloric restriction^18–20^. Attempts were made to decipher the health-promoting underpinnings of intermittent fasting^21–27^. However, the biological effects of intermittent fasting and their link to the resulting health benefits are still largely undetermined.

As detailed above, the response to fasting is dictated by considerable changes in enhancer activity and transcriptional regulation. Also, intermittent fasting produces long-term metabolic effects distinct from a single bout of fasting. Here, we aimed to examine if repeated fasting bouts are ‘remembered’ in gene expression programs, alter enhancer status and TF activity, thereby augmenting the response to future fasting events. Using a series of gene expression and enhancer accessibility profiling combined with gene knockout experiments, we found that repeated fasting bouts sensitize enhancers and gene expression programs to augment fuel production. We show that recurring fasting events are ‘remembered’ by transcriptional regulatory components, which prepare hepatocytes for the next fasting bout.

## Results

To study the effects of intermittent fasting, we designed the following experiment: 8-week-old mice were subjected to a 4-week ADF regimen in which animals had ad libitum access to food and water for 24 h followed by 24 h of access to only water (this group was termed ADF, for alternate-day fasting). In parallel, a control group of mice had unrestricted access to food and water throughout the 4 weeks (this group was termed URF, for unrestricted feeding, Fig. 1A). Food intake of ADF mice during the 4-week period was reduced by 12% compared to URF mice, showing that during the 24-h feeding period, ADF mice almost entirely compensated for the lack of food intake during the fasting day (Fig. 1B, S1A). Therefore, ADF mice are only mildly calorie restricted (as shown elsewhere^21,23,26,28,29^). In accordance, body mass was not reduced in ADF mice compared to URF mice (Fig. 1C). To more extensively compare the effects of ADF on mouse metabolism, we housed a different group of mice in metabolic phenotyping cages, placed them on a 4-week ADF regimen and measured several parameters. Food intake and body mass were comparable to values measured in conventional cages (Fig. S1B, C). Mice ran significantly higher distances on the wheel during fasting days, a known phenomenon^30^ presumably reflecting food-seeking behavior (Fig. S1D). Importantly, the respiratory exchange ratio (RER) during fasting days was 0.77 on average, showing reliance on fat oxidation. In contrast, during feeding days the RER was 0.97 on average, showing reliance on carbohydrate utilization^31^ (Fig. 1D, E). Taken together, these data show that young mice undergoing a 4-week ADF regimen show slightly reduced food intake and unaltered body mass. To maintain homeostasis, mice readily switch between fuel sources, using lipids as the principal fuel during fasting days and carbohydrates during feeding days.

**Figure 1:**
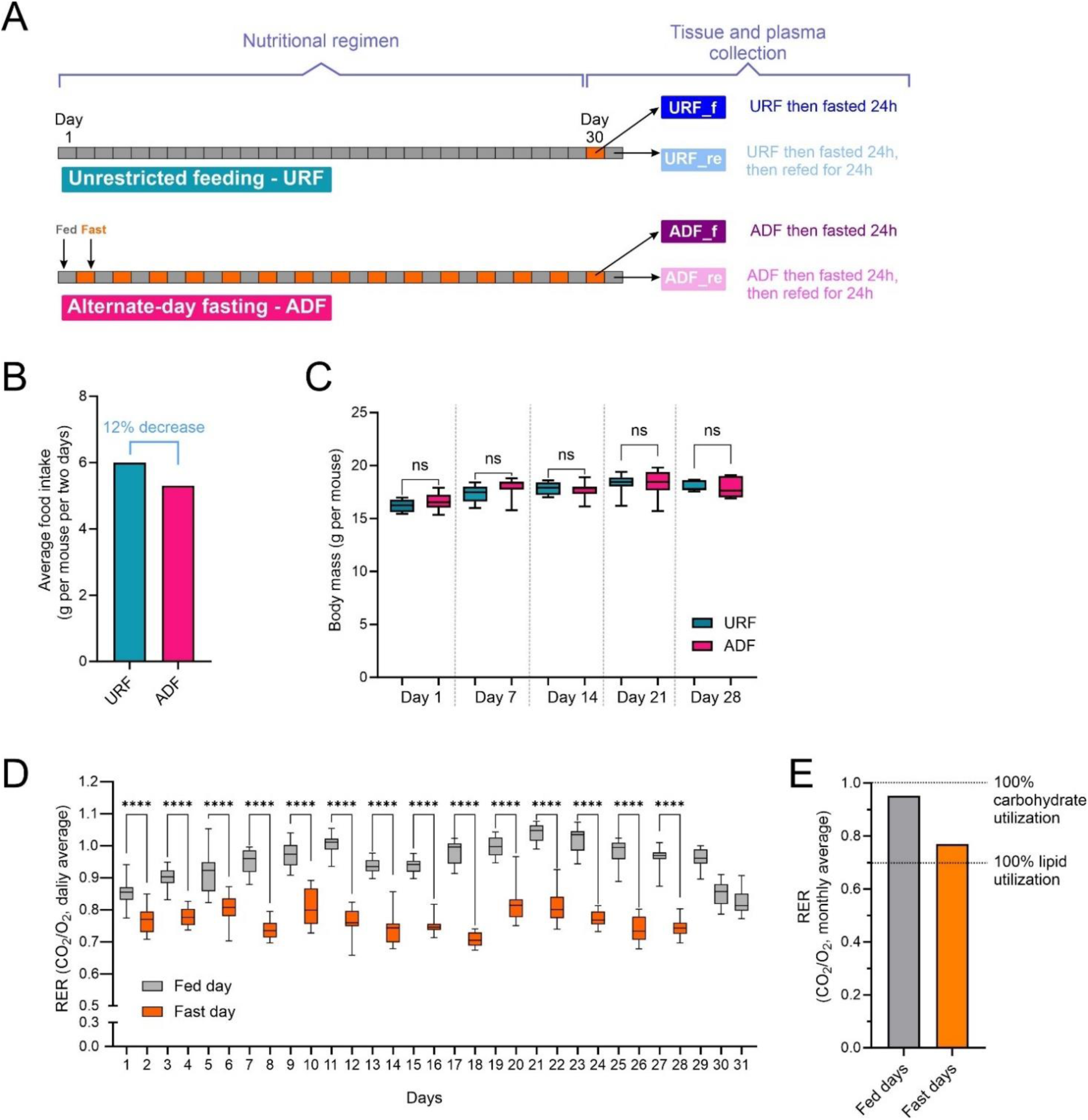
Robust fuel source switching and mildly reduced food intake maintain normal weight during ADF. **A.** Scheme of experimental design. Mice (6 per group) were put on either an unrestricted feeding (URF) or an alternate-day fasting (ADF) regimen for 30 days. Fasting periods lasted for 24 h. Livers and plasma were collected at the end of the last fasting or refeeding period. For full details, see Methods. **B.** Average food intake across the entire experiment (30 days) shows a mild decrease in food intake in the ADF group. **C.** Body mass was measured weekly, showing unchanged body mass in ADF mice as compared to URF mice. **D.** Respiratory exchange ratio (RER) was measured continuously in metabolic phenotyping cages throughout 31 days, demonstrating a switch from carbon utilization in fed days to fat utilization in fasted days. **E.** Average RER throughout the experiment duration shows preferential carbohydrate utilization on fed days and preferential lipid utilization on fast days. - Data are presented as median (C, D) or as mean (B, E). *P≤0.05, **P≤0.01, ***P≤0.001, ****P≤0.0001 by ordinary one-way ANOVA followed by Sidak post hoc analysis (C, D). Biological replicates: 5-12 (B, C) 16 (D, E).

Fuel production during fasting takes place mainly in hepatocytes where it is controlled by transcriptional and chromatin regulation^6,7^. To explore whether hepatic gene expression is differentially regulated during ADF and how ADF may affect future fasting and refeeding responses, we exposed the URF and ADF groups to acute fasting and refeeding at the end of the 4-week period. Thus, URF mice served as the control whereby half of URF mice were euthanized at the end of a 24 h fasting period (this group was termed URF_f) while the other half fasted for 24 h followed by ad libitum refeeding for 24 h at the end of which they were euthanized (this group was termed URF_re). Similarly, half of ADF mice were euthanized after 24 h of fasting (ADF_f) and the other half at the end of 24 h of refeeding (ADF_re; Fig. 1A).

We collected livers from all groups and profiled their transcriptome by RNA-seq. To deduce fasting-dependent gene regulation, differential gene expression analyses were performed between the fasted and the refed states in both URF and ADF mice. As expected, we found that fasting elicits a major transcriptional program with thousands of genes altered in both the URF and ADF groups. 2,773 genes were regulated in at least one condition, i.e. induced or repressed by fasting compared to refeeding in the URF and/or ADF groups (Table S1). Strikingly, fasting-dependent gene regulation was very different between the URF and ADF groups. Fasting led to the induction of 1,109 genes in the URF group (URF_f compared to URF_re) while only 413 genes were induced in ADF_f compared to ADF_re (Fig. 2A). Among these, 235 genes were induced in both URF_f and ADF_f. A similar trend was observed in fasting-repressed genes (Fig. 2B). This suggests that the hepatic transcriptional response to fasting is markedly affected by previous fasting events. To examine this on a more comprehensive scale, we analyzed global gene expression patterns using t-distributed stochastic neighbor embedding (t-SNE). We found that while URF_re and ADF_re cluster closely, the ADF_f and URF_f conditions are far apart, showing they meaningfully differ in their gene expression patterns (Fig. 2C). To further explore this, we compared the fold change (fasting over refeeding) of each regulated gene in both the URF and ADF groups. We plotted the fold change values of URF mice (x-axis) and ADF mice (y-axis). In a scenario where the fasting history of mice does not affect current fasting events, we would expect similar fold change values in both ADF and URF groups, resulting in a 45° angle trendline. However, we found that most genes fell below the 45° line, resulting in a 22.3° angle trendline (Fig. 2D). Thus, the general trend was that of dampened fasting-dependent gene induction in ADF, i.e. the induction level of many genes was lower in ADF compared to URF (Fig. 2D, blue-shaded area). Conversely, a smaller, albeit not negligible group of genes showed higher fasting-dependent induction in ADF (Fig. 2D, pink-shaded area). Given the obvious effect of ADF on fasting-dependent gene regulation, we directly compared gene expression between acute fasting in ‘first-time-fasters’ (URF_f) and ‘experienced fasters’ (ADF_f). We found that 1,146 genes are differentially expressed between these two conditions (Fig. S2, Table S1), aligning well with data from Fig. 2A-D showing marked transcriptional differences between URF_f and ADF_f. Taken together, these data show that the transcriptional response to fasting is dramatically different in mice that previously experienced recurrent fasting events compared to ‘first-time fasters’.

**Figure 2:**
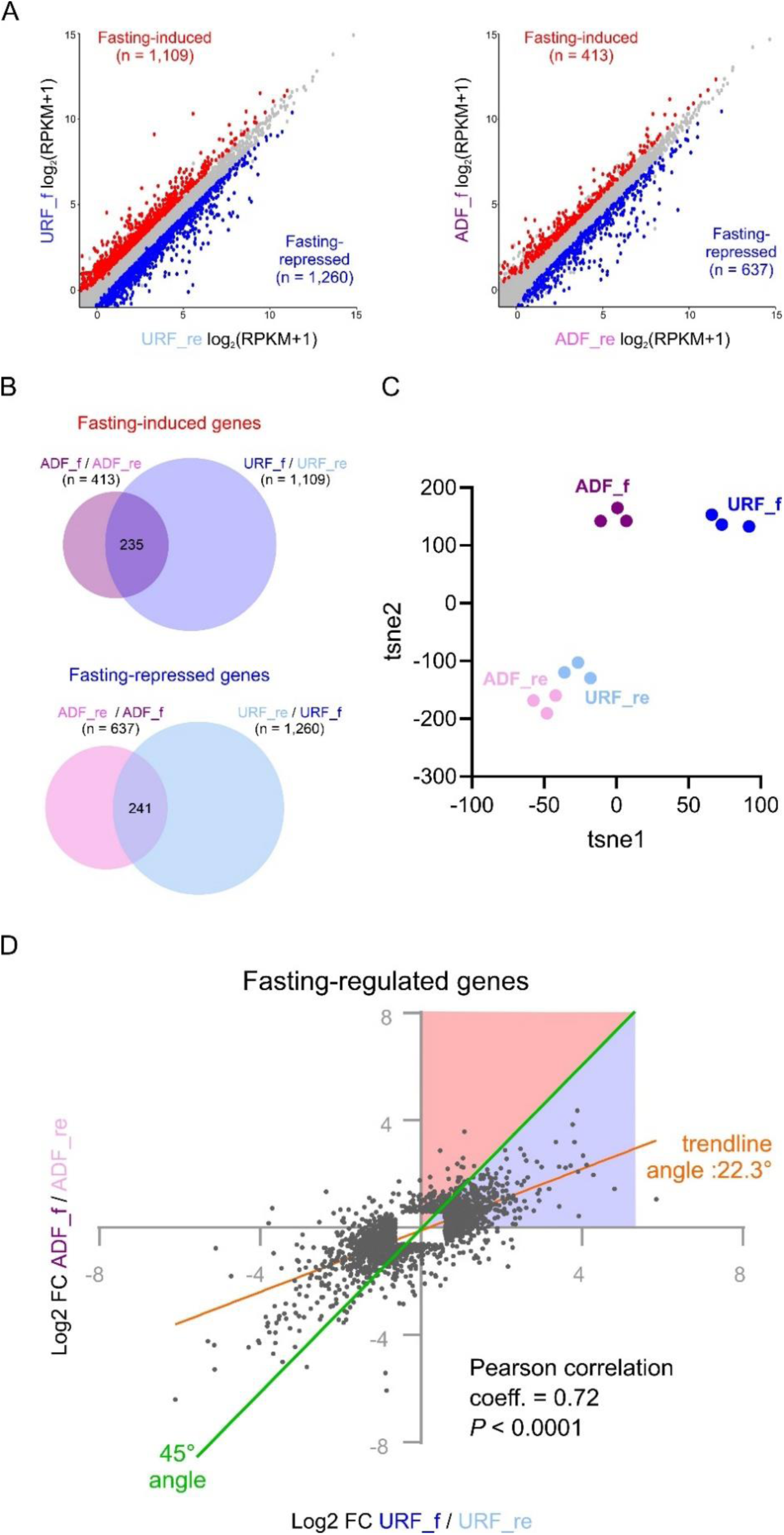
Previous fasting events drastically affect the transcriptional response to acute fasting. **A.** Normalized mRNA expression values of genes regulated by acute fasting in URF (left-hand side) and ADF (right-hand side). Red: induced genes; Blue: repressed genes; Grey: unchanged genes; Inclusion criteria for induction and repression: fold change ≥ 1.5, adj. p value ≤ 0.05; RPKM: reads per kilobase per million. **B.** Overlapping the set of genes regulated by acute fasting in URF and ADF mice shows only partial overlap, indicating that the regulatory program of gene expression during fasting is affected by past fasting events. **C.** All expressed genes in each condition were analyzed by t-distributed stochastic neighbor embedding (t-SNE) analysis, demonstrating that in the fed state URF and ADF have similar global gene expression patterns. Conversely, gene expression patterns of ADF_f are overtly different from URF_f. In all conditions, replicates cluster together, attesting to the technical quality of the data. **D.** The fold change (FC) values for all genes regulated by acute fasting are plotted; i.e. genes altered in URF_f compared to URF_re and/or genes altered in ADF_f compared to ADF_re. The x-axis value shows the URF_f over URF_re FC and the y-axis value shows the ADF_f over ADF_re FC. Comparing FC values between URF and ADF reveals two groups of genes: genes more strongly induced in ADF (pink-shaded area) and genes more weakly induced in ADF (blue-shaded area). - Data are presented as mean (A, D). Biological replicates: 3 (A-D). Genes with RPKM < 0.5 were excluded (A, C).

Next, we aimed to get a birds-eye view of gene expression patterns of *all* regulated genes across *all* four conditions. Therefore, we plotted the expression pattern of every gene regulated in at least one condition and clustered similar expression patterns (Fig. 3A). This revealed several predominant gene expression patterns represented in clusters. In line with the t-SNE analysis, two clusters showed fasting-repressed genes which were largely unaffected by ADF (Clusters 3 and 7). In contrast, we found two gene expression patterns showing a clear effect of ADF on genes induced in the fasted state as compared to the refed state (termed hereafter fasting-induced genes -FIGs). Some FIGs were less strongly induced following ADF compared to URF_f (Clusters 1, 4 and 5) while other FIGs were more strongly induced following ADF (Clusters 2 and 8).

**Figure 3:**
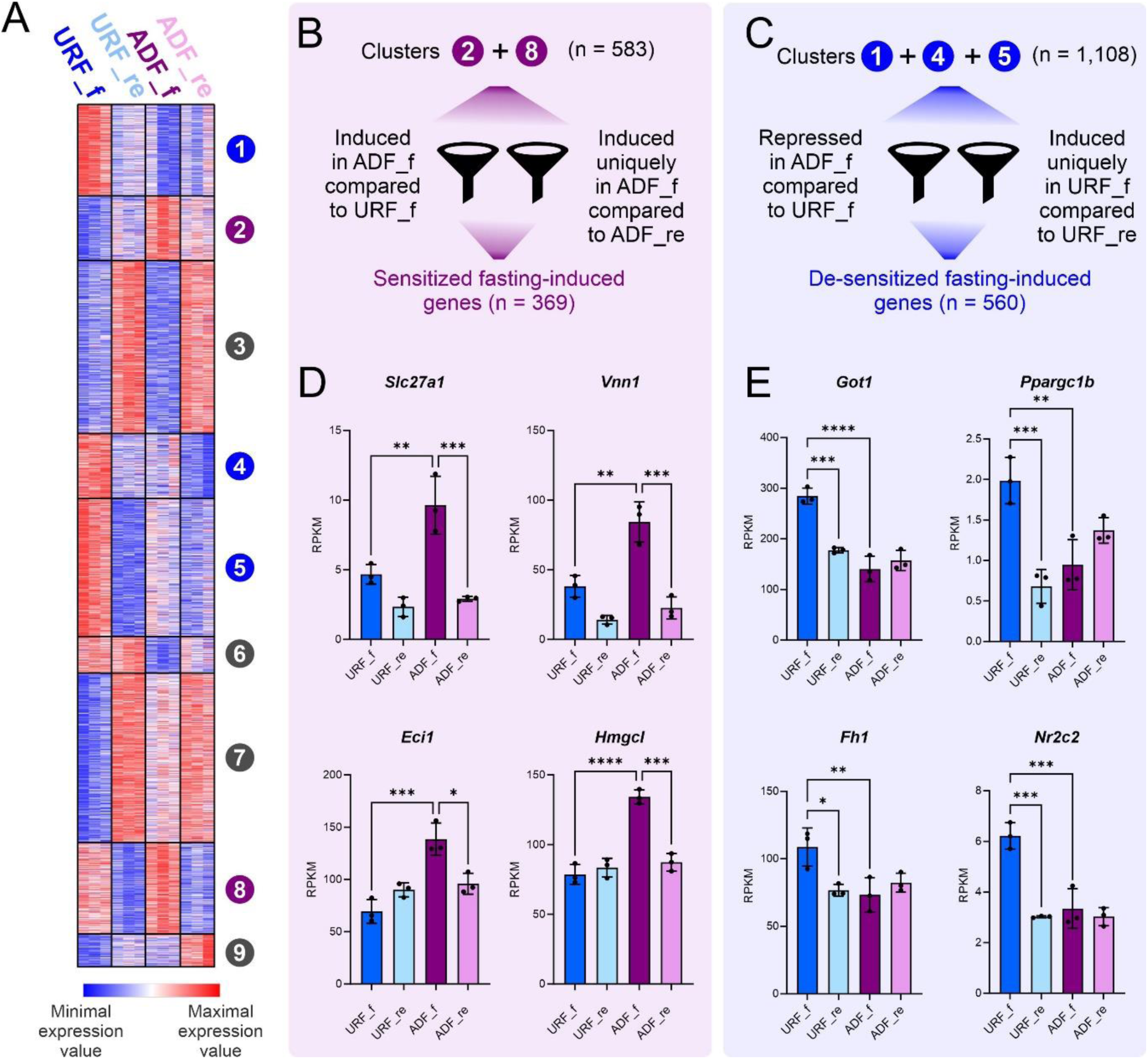
ADF sensitizes genes, augmenting ketogenic gene induction upon acute fasting. **A.** Gene clustering of all fasting-regulated genes reveals several predominant gene expression patterns. Each row represents a gene and each column represents a mouse liver sample. Notably, a group of fasting-induced genes is induced more strongly after ADF while in a different group of genes, fasting-dependent induction is dampened by ADF. Blue: minimum expression value of the gene. Red: maximum expression value of each gene (minimum and maximum values of each gene are set independently to other genes). **B, C.** Genes from relevant clusters were filtered by the specified cutoffs to strictly define sensitized and de-sensitized genes. **D.** The RPKM values of selected sensitized genes are presented, showing an increase in ADF_f compared to URF_f. **E.** The RPKM values of selected de-sensitized genes are presented, showing a decrease in ADF_f compared to URF_f. - Data are presented as mean ±S.D. (D, E). *P≤0.05, **P≤0.01, ***P≤0.001, ****P≤0.0001 by ordinary one-way ANOVA followed by Sidak post hoc analysis (D, E). Biological replicates: 3 (A-E).

Collectively, the above data (Fig. 2, S2, 3A) show that repeated fasting events sensitize the induction of certain transcriptional programs while de-sensitizing other transcriptional programs. To strictly define sensitization and de-sensitization of FIGs, we determined the following criteria. *Sensitized FIGs* (FIGs whose fasting-dependent induction is *augmented* in ADF) were defined as genes from Clusters 2 and 8 which pass at least one of the following statistical cutoffs: they are induced by fasting only in ADF but not in URF (Fig. 2A) and/or they are induced in the ADF_f condition compared to the URF_f condition (Fig. S2). *De-sensitized FIGs* (FIGs whose fasting-dependent induction is *dampened* in ADF) were defined as genes from Clusters 1, 4 and 5 which pass at least one of the following statistical cutoffs: they are induced by fasting only in URF but not in ADF (Fig. 2A) and/or they are repressed in the ADF_f condition compared to the URF_f condition (Fig. S2). This analysis revealed 369 sensitized FIGs and 560 de-sensitized FIGs (Fig. 3B, C, Table S2). Taken together, these findings reveal that repeated fasting events profoundly affect gene regulation of future fasting bouts, with some genes sensitized and others de-sensitized for future fasting-dependent induction.

To get insights as to the biological functions governed by sensitized FIGs, we searched for statistically enriched pathways and Gene Ontology terms within this group of genes. Strikingly, 7 out of 14 highly-enriched pathways were related to fatty acid oxidation, ketogenesis and the major transcriptional regulator of these processes - PPARα (Table S2). To directly quantify how many sensitized FIGs are bona fide PPARα target genes, we analyzed two published datasets that determined the PPARα-dependent hepatic gene transcriptome. The two studies determined PPARα target genes by comparing the transcriptome of wild-type mice to that of liver-specific PPARα knockout mice following either fasting- or agonist-dependent stimulation of PPARα^32,33^. We compiled a list of hepatic genes induced by PPARα (i.e., induced by fasting and/or agonist only in wild-type mice and not in PPARα knockout mice). We found that 41% (n = 153) of sensitized FIGs are PPARα target genes (Table S2). Many of these genes play a critical role in lipid catabolism and ketogenesis (e.g. *Acad, Acat, Cpt2, Hadhb, Cyp4a14, Slc27a1, Slc25a20, Hmgcl, Fgf21, Vnn1, Eci2*)^34^. Selected examples of FAO- and ketogenesis-related, PPARα-regulated FIGs sensitized by ADF are shown in Fig. 3D.

In contrast to sensitized FIGs, pathway enrichment analysis of de-sensitized FIGs did not result in an apparent pathway (selected examples of de-sensitized FIGs are shown in Fig. 3E). Interestingly, a negative regulator of PPARα (TR4, encoded by *Nr2c2*) was also de-sensitized by ADF (Fig. 3E), aligning well with the observed sensitization of PPARα target genes. Collectively, we found that the comprehensive PPARα-controlled FAO/ketogenic program is augmented in mice experiencing repeated fasting bouts.

The chromatin environment is a central factor in regulating the transcriptional response to fasting, with thousands of enhancers activated during fasting to regulate gene expression^12^. We hypothesized that sensitized and de-sensitized responses to repeated fasting are driven by alterations in enhancer activity. Therefore, we mapped accessible regions in a genome-wide manner via ATAC-seq^35^. We found a total of 182,151 accessible hepatic sites across the genome in all conditions. Previous data suggests most accessible regions in the genome are cis-regulatory regions (in particular promoters or enhancers)^36^. Only 6.5% of hepatic accessible sites were promoter-proximal regions (Fig. S3A), suggesting that the majority of accessible sites we found are enhancers. Accordingly, the motifs enriched in accessible hepatic sites are motifs associated with hepatic enhancers such as C/EBP, HNF4α and FoxA (Fig. S3B)^37^.

To evaluate the *dynamics* in enhancer accessibility imposed by ADF, we measured differential accessibility between the ADF_f and URF_f groups. Remarkably, although both groups were collected after fasting, their enhancer landscape was considerably different with 23,629 sites changing their accessibility due to ADF. The enhancers showing altered accessibility were roughly evenly divided between sensitized enhancers (showing increased accessibility in ADF_f compared to URF_f, n = 11,571) and de-sensitized enhancers (decreased in ADF_f, n = 12,058, Fig. 4A, Table S3). This demonstrates that repeated fasting events lead to vast changes in chromatin organization with 13% of liver accessible sites showing altered accessibility following ADF.

**Figure 4:**
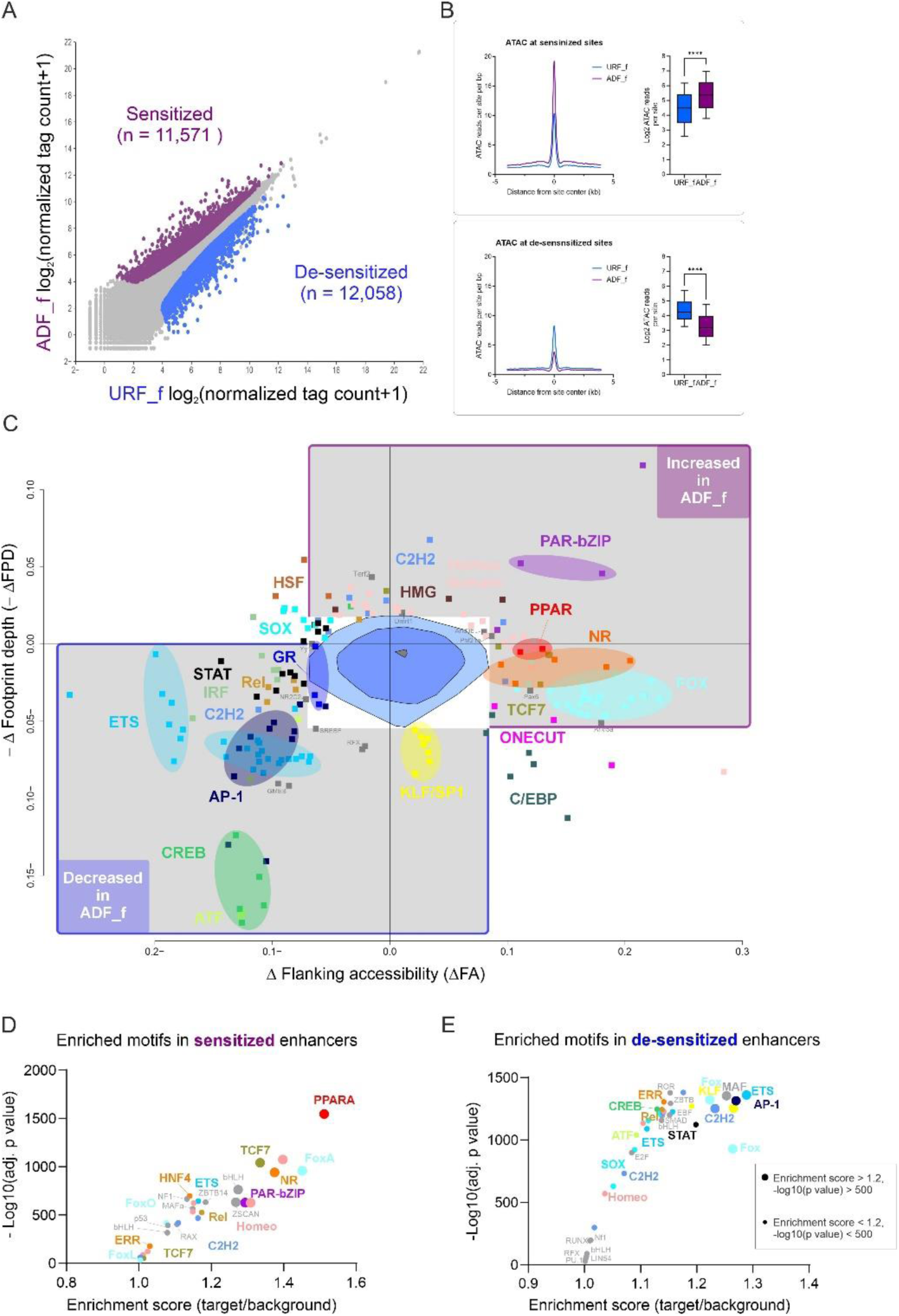
Enhancer accessibility near ketogenic genes is sensitized by ADF. **A.** ATAC accessible sites with differential accessibility in ADF_f as compared to URF_f are shown. Unchanged sites are in grey, induced sites are in purple, repressed sites are in blue. Inclusion criteria for induction and repression: fold change ≥ 1.5, adj. p value ≤ 0.05. **B.** Quantification of chromatin accessibility at sensitized and de-sensitized enhancers. **C.** Bivariate Genomic Footprinting (BaGFoot) analysis reveals TFs predicted to have increased (top right region) or decreased (bottom left region) activity in ADF_f compared to URF_f. TFs with a similar or identical DNA binding motif were marked with the same color. **D.** Motif enrichment analysis shows TFs whose motifs are enriched in sensitized enhancers. **E.** Motif enrichment analysis shows TFs whose motifs are enriched in de-sensitized enhancers. - Data are presented as log2 normalized mean tag counts (A). P≤0.05, **P≤0.01, ***P≤0.001, ****P≤0.0001 by unpaired two-tailed t-test (B). Biological replicates: 3 (A, B, C). Values with RPKM<0.5 were omitted (A). Only motifs with -log10(p-value) > 10 are shown (D, E).

Such a widespread effect on enhancer accessibility is expected to be caused by profound changes in TF activity that in turn activate enhancers, leading to increased accessibility. We sought to uncover the TFs driving this effect on chromatin. Thus, we analyzed accessible sites in two independent unbiased approaches. First, we used BaGFoot, a tool that predicts TF activity from TF footprints as well as chromatin accessibility changes^13^. BaGFoot detects all known TF motif occurrences across all accessible regions. Then, it quantifies both the accessibility flanking the motif (termed ‘flanking accessibility’) and the footprint depth within the motif. An increase in footprint depth and/or flanking accessibility suggests the TF is more active in the tested condition on a genome-wide scale^13,38^. Using BaGFoot, we compared the ADF_f and URF_f conditions for changes in flanking accessibility and footprint depth across all accessible regions. We found multiple TFs with altered activity between conditions (Fig. 4C), TFs with increased activity in ADF_f appear in the top right area and TFs with decreased activity in ADF_f appear in the bottom left area. The most prominent TFs with increased activity in ADF_f were PPARα, PAR bZIP TFs, Fox proteins and several nuclear receptors. The predominant TFs with increased activity in URF_f were CREB, AP-1 and ETS proteins. Thus, BaGFoot revealed several TFs associated with sensitized and de-sensitized enhancers. In addition to BaGFoot, we directly analyzed the groups of sensitized and de-sensitized enhancers (Fig. 4A) to find significantly enriched TF motifs in each population. Motif enrichment analysis was highly concordant with BaGFoot results with PPARα, PAR bZIPs and Fox proteins highly enriched in sensitized enhancers (Fig. 4D). Motifs enriched in de-sensitized enhancers were also concordant with BaGFoot results with AP-1 and ETS motifs highly enriched and the CREB motif also among the top enriched motifs (Fig. 4E).

The evidence from both gene expression (Fig. 3, Table S2) and chromatin data (Fig. 4C, D) clearly suggested a role for PPARα in enhancer sensitization. We examined the possibility that this is mediated via an ADF-dependent increase in PPARα protein levels. However, total PPARα protein levels were unchanged in ADF (Fig. S4A). Thus, we examined the possibility of preferred PPARα binding at sensitized enhancers. We performed chromatin immunoprecipitation sequencing (ChIP-seq) for PPARα in URF_f and ADF_f. Most PPARα binding sites were located in promoter-distal regions (89%, Fig. S4B) and the PPARα response element was the top enriched motif among PPARα binding site, as expected (Table S4). Strikingly, we found that PPARα occupancy was enriched in sensitized enhancers compared to de-sensitized enhancers (Fig. 5A, B). Moreover, PPARα occupancy in sensitized enhancers was significantly higher in the ADF_f condition compared to URF_f (Fig. 5C). PPARα occupancy was also increased near sensitized genes in the ADF_f condition as compared to the URF_f condition (Fig. 5D). Therefore, even though we compared two groups in which mice were fasted for a period known to potently increase PPARα activity and binding, ADF mice show heightened PPARα activity. This aligns with the increased PPARα BaGFoot signal, PPARα motif enrichment and augmented PPARα-related gene expression in ‘experienced fasters’ (ADF_f). Indeed, the loci of ADF-sensitized PPARα target genes show enhancer sensitization as well as increased PPARα occupancy following ADF_f (compared to URF_f, Fig. 5E). Taken together, these data show that a ketogenic gene program is increased in ADF, together with increased accessibility of enhancers and heightened binding of the major ketogenic TF – PPARα.

**Figure 5:**
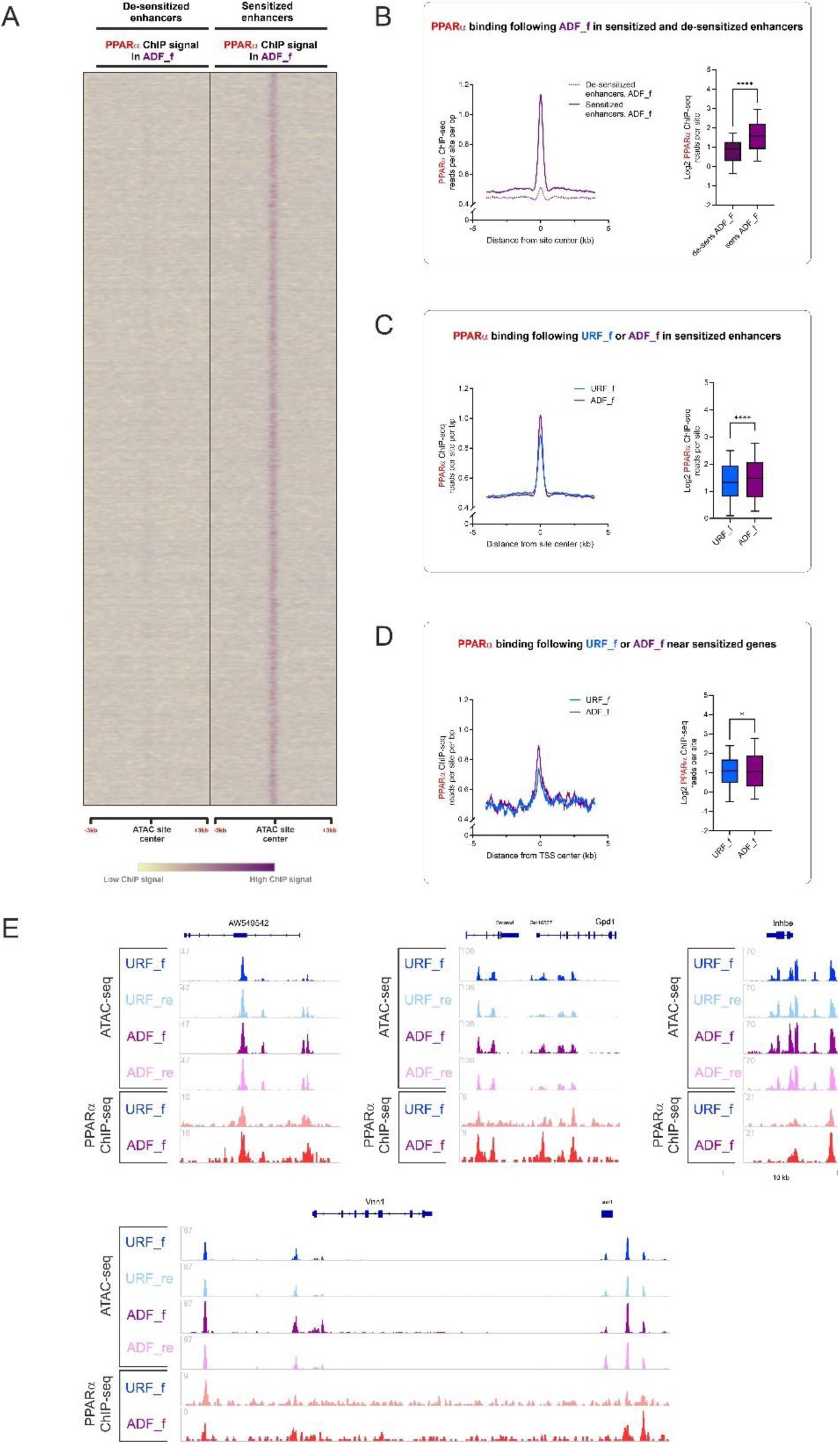
PPARα preferentially binds sensitized enhancers following ADF. **A, B.** Quantification of PPARα occupancy at sensitized and de-sensitized enhancers shows PPARα preferentially binds sensitized enhancers with minimal occupancy at de-sensitized enhancers. **C.** Quantification of PPARα occupancy at sensitized enhancers in either the ADF_f or URF_f states shows higher PPARα occupancy following ADF_f. **D.** Quantification of PPARα occupancy near sensitized genes in either the ADF_f or URF_f states shows higher PPARα occupancy following ADF_f. **E**. Genome browser images of selected sensitized enhancers proximal to sensitized PPARα target genes show enhancer sensitization and increased PPARα binding following ADF. - P≤0.05, **P≤0.01, ***P≤0.001, ****P≤0.0001 by unpaired two-tailed t-test (B, C, D). Biological replicates: 2 (A-D). Representative replicate is presented (E).

While accessibility is a major determinant of enhancers, other factors also indicate enhancer activity. These include acetylation of lysine 27 in Histone H3 (H3K27ac) and occupancy of p300^9^. Additionally, in liver, the binding of the lineage-determining TFs FoxA1/2, HNF4α and C/EBPβ is also indicative of enhancer activity as these TFs facilitate the binding of other TFs^37^. These factors are often termed pioneer TFs as they are the first to bind enhancers, leading to their increased accessibility and activity^39^. To evaluate these enhancer markers in sensitized enhancers, we quantified ChIP-seq signal strength of each marker in sensitized and de-sensitized enhancers. We found that the occupancy of p300 and liver pioneer factors is substantially heightened in sensitized enhancers compared to de-sensitized enhancers (Fig. S4C). We then stratified sensitized enhancers into two classes: sensitized enhancers with or without PPARα binding. Sensitized enhancers harboring PPARα binding sites were dramatically more enriched with pioneer factor binding and enhancer marks (Fig. S4C). These data suggest that PPARα-bound sensitized enhancers are pre-bound with liver pioneer factors and are enriched with enhancer markers, making them focal points with strong enhancer characteristics.

To evaluate the effects of ADF outside the liver, we measured several circulating parameters. Plasma triglycerides and high-density lipoprotein (HDL) levels were unchanged between groups (Fig. S5A, B). Low-density lipoprotein (LDL) levels were decreased (Fig. S5C) in ADF_f, aligning with LDL lowering in humans undergoing intermittent fasting^17^. The levels of BHB, the predominant ketone body, were elevated following fasting in both URF_f and ADF_f as compared to their refed counterparts. Notably, the fasting-dependent increase in ketogenesis was significantly more pronounced in ADF mice as BHB levels were higher in ADF_f compared to URF_f (Fig. 6A). The augmented production of BHB in ADF mice is in complete agreement with the observed increases in FAO/ketogenic gene program, enhancer sensitization and increased PPARα binding at sensitized enhancers following ADF. In contrast to ketone bodies, fasting blood glucose levels were unaffected by ADF (Fig. S5D).

**Figure 6:**
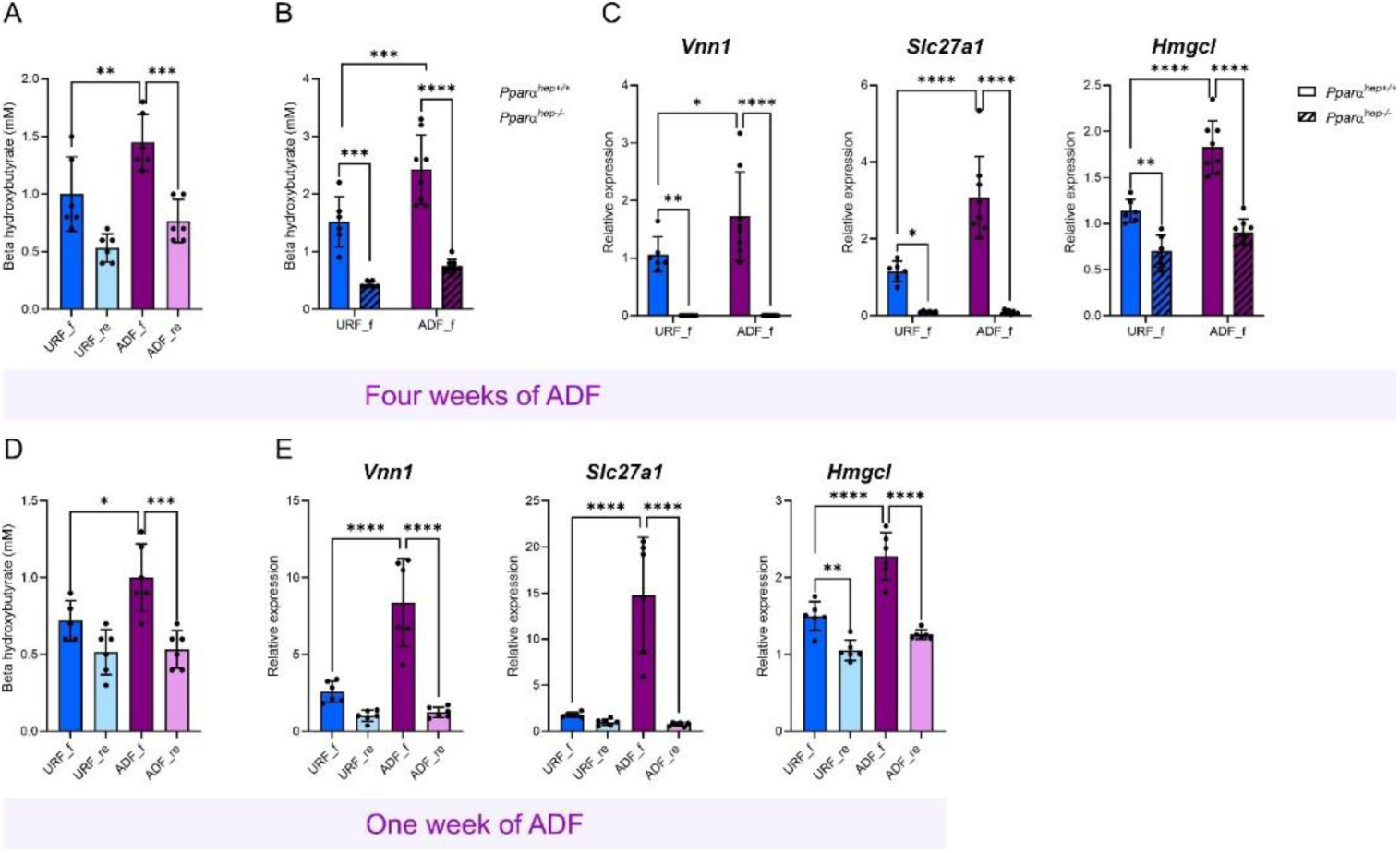
The levels of BHB are augmented following ADF. **A.** The plasma levels of BHB were measured after 4 weeks of ADF, showing an increase in BHB in ADF_f as compared to URF_f. **B.** The plasma levels of BHB were measured after 4 weeks of ADF, showing an increase in BHB in ADF_f as compared to URF_f only in Pparα^hep+/+^ mice and not in Pparα^hep-/-^ mice. **C.** The mRNA levels of selected sensitized genes were examined by qPCR after 4 weeks of ADF in, showing sensitization only in Pparα^hep+/+^ mice and not in Pparα^hep-/-^ mice. **D.** The plasma levels of BHB were measured after 1 week of ADF, showing an increase in BHB in ADF_f as compared to URF_f. **E.** The mRNA levels of selected sensitized genes were examined by qPCR after 1 week of ADF. - Data are presented as mean ±S.D. P≤0.05, **P≤0.01, ***P≤0.001, ****P≤0.0001 by ordinary one-way ANOVA followed by Sidak post hoc analysis. Biological replicates: 6-8.

The results in Figures 4 and 5 strongly suggest that sensitization is governed by PPARα. To causally link PPARα to sensitization and augmented ketogenesis in ADF, we employed hepatocyte-specific PPARα deficient mice (PPARα^hep-\-^). Similarly to WT mice, ADF in PPARα^hep-\-^ mice did not affect body mass (Fig. S5E) nor fasting glucose levels (Fig. S5F). While BHB levels were increased by ADF in PPARα^hep+/+^ mice, BHB levels were significantly impaired in PPARα^hep-/-^ mice and were not augmented by ADF (Fig. 6B). Accordingly, sensitization of genes functioning in lipid catabolism and ketogenesis was abolished in PPARα^hep-/-^ mice (Fig. 6C).

We were interested to evaluate whether the ADF-dependent changes we observed require 4 weeks of an ADF regimen or whether a shorter regimen will suffice. Similarly to BHB levels after 4 weeks of ADF, we found that BHB levels were also increased by ADF after only 1 week of ADF (Fig. 6D), as was gene sensitization (Fig. 6E). Collectively, these data show that ADF sensitizes the liver to produce more BHB through a PPARα-governed transcriptional program.

Taken together, we show that PPARα-bound enhancers regulating FAO/ketogenesis are sensitized by repeated fasting events. Accordingly, the ketogenic transcriptional program is sensitized by ADF and the plasma levels of BHB are augmented.

## Discussion

Organisms are often exposed to recurring environmental signals such as light/darkness, seasonal rhythms, cold/heat, scarcity/availability of food, etc. The response to many of these signals is brought about through regulating gene transcription. This raises two intriguing questions: **(a)** Do mammals adapt to frequently-encountered challenges by a cellular memory mechanism? **(b)** Do recurrent signals sensitize transcriptional programs to maximize future responses and increase survival?

Mammals evolved to maintain homeostasis during frequent and prolonged fasting periods. Dietary regimens that include repeated fasting bouts such as ADF are becoming increasingly popular as studies have shown their outstanding health benefits. Our study, as well as many others, show no change in body mass and little-to-no change in food intake during ADF^23,24,26,28,29^. These findings preclude the possibility that the benefits of ADF are a result of reduced body mass or calorie restriction but are rather an effect of fasting per se. Therefore, while the benefits of ADF are well described, how recurrent fasting events promote them is unclear. To better understand this, we set out to decipher how ADF affects the liver’s response to future fasting events.

We found that ADF profoundly changes the transcriptional program and chromatin landscape of the liver to support a robust ketogenic program. Mice that have experienced several fasting bouts respond in a profoundly different manner to a following acute fast when compared to ‘first-time fasters’. The hepatic chromatin landscape of ‘experienced fasters’ is altered with many enhancers being sensitized for activation by previous fasting events. In turn, the induction of ketogenic genes is sensitized and the production of ketone bodies is augmented. Thus, by sensitizing enhancers and priming them for activation, ADF leads to a more robust fasting response in the following fasting bouts.

The transcriptional response to fasting was considerably affected by previous fasting events with some fasting-induced genes being de-sensitized (i.e. their induction was dampened following ADF) while others sensitized by ADF (i.e. their induction was augmented following ADF). What we term here as ‘gene sensitization’ is reminiscent of the coined term ‘transcriptional memory’ which was studied mostly in non-mammalian models or cultured cells. In the transcriptional memory model, a cell is able to mount a more robust transcriptional response to a signal it has previously encountered^40–42^. While there are certain similarities between transcriptional memory and our observation of gene sensitization, there are important differences: First, we describe here a mammalian response to a recurring and prevalent nutritional state. Second, we describe a bifurcated effect whereby some fasting-induced genes are sensitized while others are de-sensitized. Third, transcriptional memory was not reported to be associated with enhancers but rather driven by other factors.

The group of sensitized genes was highly enriched with genes responsible for lipid catabolism and ketogenesis. Accordingly, BHB levels were increased in fasted mice following ADF. A previous study reported variable levels of ketone bodies in ADF_f that were either reduced or unchanged when compared to URF_f ^25^. The difference between our observations and those of Li et al. might stem from a different experimental setup (for example, we used female mice while Li et al. used male mice). Nevertheless, we observed a reproducible increase in BHB following 1 and 4 weeks of ADF. Thus, our findings suggest that recurring fasting rewires the hepatic fasting response to augment lipid catabolism and ketogenesis. Indeed, a study in humans showed increased ketonemia after 4 weeks of ADF^43^. The health benefits of ADF include an improved lipid profile. It is therefore tempting to speculate that some of the health benefits of ADF are due to a shift toward lipid catabolism.

We found a major effect of ADF on the enhancer landscape with thousands of enhancers either sensitized or de-sensitized. Sensitized enhancers were associated with a FAO/ketogenic gene program. This shows that previous fasting events ‘prime’ ketogenic-related enhancers, leading to their increased activity in the next fasting bout. Several pieces of evidence link the transcription factor PPARα to enhancer sensitization: a) its motif is enriched within sensitized enhancers; b) the accessibility flanking the motif is increased following ADF; c) PPARα occupancy is increased in sensitized enhancers following ADF. Importantly, gene sensitization and augmented ketonemia is ablated in hepatocyte-specific PPARα-deficient mice. In addition, one of the genes de-sensitized by ADF is *Nr2c2*. This gene encodes TR4, a factor known to antagonize PPARα activity^44^. Thus, the increased activity of PPARα in ADF may be supported by lowered TR4 expression.

In summary, we have shown here that repeated fasting bouts sensitize enhancers and gene induction to produce more robust ketogenesis. We showed that these effects are evident as early as 1 week after the commencement of the regimen, suggesting sensitization is a relatively quick response. We show that recurring fasting events are ‘remembered’ by transcriptional regulatory components, which prepare hepatocytes for the next fasting bout. Considering that, in addition to repeated fasting, organisms are routinely exposed to other recurring environmental signals, our findings may shed light on transcriptional regulation as a mediator of cellular adaptation to repeated signals and physiological habituation to the environment.

## Methods

### Animals

Female, 6 weeks-old mice (C57BL/6J) were randomly assigned to either the URF or ADF groups (12 mice per group). The experiment started after 1 week of acclimation. The URF group had ad libitum access to food (Teklad TD2018) and water for 30 days. The ADF group had ad libitum access to food and water for 24 h followed by ad libitum access to only water for 24 h. The ADF group went through 15 cycles of fasting-refeeding (a total of 30 days). Food was removed at the beginning of the inactive phase (shortly after lights on, zeitgeber time 1). Food was put back in the cage 24 h later. On day 31 all mice in the experiment underwent 24 h of fasting. Half of the mice were euthanized at the end of the fasting period (6 mice from the URF group and 6 mice from the ADF group). The remaining mice were refed and euthanized 24 h after the food was put back in the cage (6 mice from the URF group and 6 mice from the ADF group). At the end of the experiment mice were anesthetized and euthanized (ketamine:xylazine 30:6 mg/ml), the liver was excised and blood was collected into heparin-coated tubes to allow plasma isolation.

For the metabolic cages experiment, 16 mice (female, 6 weeks-old mice, C57BL/6JOlaHsd) were singly housed in metabolic phenotyping cages (Promethion Core, Sable Systems). The experiment started after 1 week of acclimation. The ADF experimental design described above was replicated with all mice undergoing ADF for 30 days. In metabolic cages, access to food was controlled remotely by programmed opening and closing of the food access control door.

Pparα hepatocyte-specific knockout (Pparα^hep-/-^) mice were generated at INRAE’s rodent facility (Toulouse, France) by mating the floxed-Pparα mouse strain with C57BL/6J albumin-Cre transgenic mice, as described previously^45^, to obtain albumin-Cre^+/-^Pparαflox/flox mice. Albumin-Cre^-/-^Pparα^flox/flox^ (Pparα^hep+/+^) littermates were used as controls. Genotyping was performed using an established protocol^45^. Fourteen 18-week-old females Pparα^hep+/+^ and fourteen 18-week-old females Pparα^hep-/-^ were randomly assigned to either the URF (6 mice per genotype) or ADF groups (8 mice per genotype). The experiment started after 1 week of acclimation. The URF group had ad libitum access to food (Safe 04 U8220G10R) and water for 30 days. The ADF group had ad libitum access to food and water for 24 h followed by ad libitum access to only water for 24 h. The ADF group went through 15 cycles of fasting-refeeding (a total of 30 days). Food was removed at the beginning of the inactive phase (shortly after lights on, zeitgeber time 1). Food was put back in the cage 24 h later. On day 31 all mice in the experiment underwent 24 h of fasting and were euthanized at the end of the fasting period. Following sacrifice by cervical dislocation, the liver was removed, weighed, snap frozen in liquid nitrogen, and stored at -80 °C until use.

All animal procedures are compatible with the standards for the care and use of laboratory animals. The research has been approved by the Hebrew University of Jerusalem Institutional Animal Care and Use Committee (IACUC). The Hebrew University of Jerusalem is accredited by the NIH and by AAALAC to perform experiments on laboratory animals (NIH approval number: OPRR-A01-5011). The Pparα^hep-/-^ experiments were approved by an independent ethics committee under the authorization number 45717-2023111017412475.

### RNA preparation, reverse transcription and quantitative PCR (qPCR)

Total RNA was isolated from liver pieces (30 mg) using a NucleoSpin kit (Macherey-Nagel cat# 740955.25) according to the manufacturer’s protocol. For qPCR, 1 μg of total RNA was reverse transcribed to generate cDNA (Quantabio cat# 76047-074). qPCR was performed using C1000 Touch thermal cycler CFX96 and CFX Opus 384 instruments (Bio-Rad) using SYBR Green (Quantabio cat# 101414-276). Gene values were normalized with housekeeping genes (*Gapdh*). The sequences of primers used in this study are:

*Gapdh* - Fwd: CAGGAGAGTGTTTCCTCGTCC, Rev: TTTGCCGTGAGTGGAGTCAT

*Hmgcl* - Fwd: ACTACCCAGTCCTGACTCCAA, Rev: TAGAGCAGTTCGCGTTCTTCC

*Vnn1* - Fwd: CGCACCTGTGGTAGTTCAGT, Rev: GGTTAACACAGGTCCCGAGG

*Slc27a1* - Fwd: CAGAACTTCCCAGTCCAGACTTC, Rev: ACGTACACACAGAACGCCG

### Western Blot

RIPA buffer (50mM Tris-HCl, 150mM NaCl, 1% triton, 0.5% sodium deoxycholate, 0.1% SDS) with protease inhibitors (Sigma, cat# P2714) was added to liver pieces (70 mg) followed by 1 min of homogenation (Bead Ruptor 12, Omni international) and centrifugation. Protein samples (50 µg) were loaded on 12% polyacrylamide SDS gels. Proteins were transferred (Trans-Blot Turbo, Bio-Rad; cat# 1704158) to a nitrocellulose membrane (Trans-Blot Turbo Transfer Pack, Bio-Rad; cat# 1704158), blocked for 1 h with 5% low-fat milk, and incubated for 16 h with primary antibody (PPARα, Santa Cruz Biotechnology cat# sc-398394; histone H3, Cell Signaling Technologies cat# 14269) diluted 1:1000 in tris-buffered saline (0.5% Tween, 5% bovine serum albumin). Membranes were incubated with secondary peroxidase AffiniPure goat anti-mouse (1:10,000, Jackson Laboratory; cat# 115-035-146) for 1 h, followed by washes and a 1-minute incubation with western blotting detection reagent (Cytiva Amersham ECL prime, cat# RPN2232). Imaging and quantification were done with ChemiDoc (Bio-Rad).

### Blood measurements

Plasma samples were analyzed for triglycerides, cholesterol, LDL and HDL using a Cobas c111 (Roche Diagnostics) automated clinical chemistry analyzer that was calibrated according to manufacturer guidelines. Glucose and BHB were measured directly on blood with FreeStyle Optium Neo and glucose strips (Abbott cat# 7069470) or beta ketone test strips (Abbott, cat# 7074270).

### Chromatin immunoprecipitation (ChIP)

ChIP was performed as previously described^46^ with modifications: Liver pieces (150 mg) were cross-linked with phosphate-buffered saline (PBS) containing 2mM disuccinimidyl glutarate (DSG, Santa Cruz Biotechnology, cat# sc-285455). Livers were homogenized with a Dounce homogenizer and rotated for 30 min at room temperature. Samples were centrifuged and the pellet was resuspended with PBS containing 1% formaldehyde (Electron Microscopy Sciences, cat# 15714) for further crosslinking. After 10 min, samples were quenched with 0.125 M glycine for 5 min. Samples were then centrifuged, washed with PBS, resuspended in ChIP lysis buffer (0.5% SDS, 10mM EDTA, 50mM Tris-HCl pH8) and sonicated (Bioruptor Plus, Diagenode) to release 100–1000 bp fragments. Samples were diluted 1:5 with ChIP dilution buffer (170 mM NaCl, 17 mM Tris-HCl pH8, 1.2 mM EDTA, 1.1% Triton x-100, 0.01% SDS).

Antibody against PPARα (Merck Millipore MAB3890, 4 µg per sample) was conjugated to magnetic beads (Sera-Mag, Merck, cat# GE17152104010150) for 2 h at 4°C. Chromatin was immunoprecipitated with antibody-bead conjugates for 16 h at 4°C. Immunocomplexes were washed sequentially with the following buffers: low salt buffer (0.01% SDS, 1% Triton x-100, 2 mM EDTA, 20 mM Tris-HCl pH 8, 150 mM NaCl), high salt buffer (0.01% SDS, 1% Triton x-100, 2 mM EDTA, 20 mM Tris-HCl pH 8, 500 mM NaCl), LiCl buffer (0.25M LiCl, 1% IGEPAL CA630, 1% deoxycholic acid, 1mM EDTA, 10 mM Tris pH 8.1) and twice with TE buffer (10 mM Tris-HCl, 1 mM EDTA pH 8). Chromatin was eluted, de-crosslinked for 4 h at 65°C and deproteinized with proteinase K (Hy Labs, cat# EPR9016) for 1 h at 50°C. DNA was subsequently isolated using MinElute DNA purification kit (Qiagen cat# 20-28006).

### RNA-seq

For quality control of RNA yield and library synthesis products, the RNA ScreenTape and D1000 ScreenTape kits (both from Agilent Technologies), Qubit RNA HS Assay kit, and Qubit DNA HS Assay kit (both from Invitrogen) were used for each specific step. mRNA libraries were prepared from 1 µg RNA using the KAPA Stranded mRNA-Seq Kit, with mRNA Capture Beads (KAPA biosystems, cat# KK8421). The multiplex sample pool (1.6 pM including PhiX 1%) was loaded on NextSeq 500/550 High Output v2 kit (75 cycles) cartridge, and loaded onto the NextSeq 500 System (Illumina), with 75 cycles and single-read sequencing conditions.

### ATAC-seq

ATAC-seq was performed as described in our published protocol^47^. Briefly, nuclei were isolated using a hypotonic buffer and Dounce homogenizer. Nuclei were tagmented using Tn5 transposase loaded with Illumina adapters. Tagmented DNA was PCR-amplified with sample-specific indices. The resulting library was size-selected to DNA fragments of 150-800 nt. The multiplex sample pool (1.6 pM including PhiX 1%) was loaded on NextSeq 500/550 High Output v2 kit (75 cycles) cartridge with paired-read sequencing conditions. Each sample was sequenced at a depth of at least 5 X 10^7^ reads.

### Sequencing data analyses

Fastq files were mapped to the mm10 mouse genome assembly using Bowtie2^48^ with default parameters. Tag directories were made using the makeTagDirectory option in the HOMER suite^49^. Selected gene loci were visualized by the integrated genome browser (IGV)^50^.

### Differential gene expression

Differential gene expression was evaluated by DESeq2^51^ via the HOMER suite under default parameters. Genes were determined as differentially expressed between two conditions if they pass these cutoffs: fold change ≥ 1.5, adjusted p-value ≤ 0.05.

### t-distributed stochastic neighbor embedding (t-SNE)

performed by using the Rtsne package (R version 4.2.1). The analyzed values are log2(RPKM+1). Genes with RPKM < 0.5 were excluded.

### k-means clustering

all genes regulated in at least one treatment (fold change ≥ or ≤ 1.5, adj. p value ≤ 0.05) were included in the analysis. The normalized tag counts of each gene were used for the clustering analysis. The normalized tag counts of each gene were used for the analyses. Morpheus (https://software.broadinstitute.org/morpheus) was used to cluster genes under these parameters: k = 9; metric - one minus Pearson correlation; maximum iterations – 1000. Blue - minimum value of the gene; Red – maximum value of each gene (minimum and maximum values of each gene are set independently to other genes).

### ATAC-seq analyses

Peak-calling was performed by MACS2 (narrowPeak option)^52^. Differential enhancer activity was measured by DEseq2 (fold change ≥ 1.5, adj. p value ≤ 0.05). Normalized tag counts were used to visualize DESeq2 results in a scatter plot. Genomic annotations were made by the HOMER suite (annotatePeaks option, parameter -annStats).

### Bivariate genomic footprinting (BaGFoot)

BaGFoot was performed as described ^13^. Briefly, the three replicates from each condition (URF_f, ADF_f) were merged into a single BAM file. Accessible sites were called for each BAM file using MACS2. The footprint depth (FPD) and flanking accessibility (FA) were calculated for each known motif across all accessible sites. The difference (Δ) between ADF_f and URF_f were calculated and plotted on the bag plot.

### De novo motif enrichment analysis

To unbiasedly detect enriched motifs, we performed a de novo motif enrichment analysis using the findMotifsGenome option in HOMER (parameter -size given). The entire enhancer landscape (all ATAC accessible sites across all conditions) was used as background to account for possible sequence bias. Using the entire enhancer landscape as background ensures that prevalent motifs appearing across liver enhancers will not be falsely detected as specifically enriched in the examined subset of enhancers. In motif enrichment analyses of total ATAC accessible sites and PPARα binding sites, the background was automatically selected by HOMER to account for GC bias and other sequence biases.

### Aggregate plots and box plots

Tag density of ATAC or ChIP signal around ATAC site center or transcription start site (TSS) were analyzed using the HOMER suite. In aggregate plots, the tag count (averaged across all sites) per site per bp was calculated using the HOMER suite (annotatePeaks, option -size 8000 -hist 10). In box plots, tag count +/- 200 bp around the site center (averaged across all sites) was calculated using the HOMER suite (annotatePeaks, option -size 400 -noann). In both aggregate plots and box plots, the data is an average of all available replicates. In all box plots, the 10-90 percentiles are plotted.

### Pathway enrichment analysis

Performed by GeneAnalytics^53^.

### Analysis of data from the literature

PPARα target genes in hepatocytes were found from analyzing previously published data^32,33^. Differential gene expression was evaluated by DESeq2 via the HOMER suite under default parameters. Genes were determined as differentially expressed between two conditions if they pass these cutoffs: fold change ≥ 1.5, adjusted p-value ≤ 0.05. ChIP-seq data for enhancer markers and liver pioneer factors were analyzed from published datasets: H3K27ac^12^, p300^54^, Foxa1/2^55^, HNF4α^37^, C/EBPβ^12^.

## Materials availability

This study did not generate new unique reagents.

## Acknowledgments

We would like to thank Dr. Walter Wahli and Dr. Alexandra Montagner for providing key reagents and advice; Dr. Ofer Gover for help in metabolic phenotyping cages experiments; Dr. Asaf Marco and Dr. Ofri Karmon for help in experiments; Dr. Idit Shiff for support in high throughput sequencing and to Michael Fadi Saikali for help in R analyses. Graphical abstract created by HandMed - Dr. Yitzchak Yadegari.

## Author Contributions

Contribution categories were adopted from CRediT (Contributor Roles Taxonomy)

N.K Conceptualization, Methodology, Visualization, Software, Validation, Formal analysis, Investigation

M. C-N Conceptualization, Methodology, Software, Validation, Formal analysis, Investigation

J. B Investigation, Validation and Formal analysis

D. G. Investigation, Visualization, Formal analysis

D. M-A Investigation and Validation

D. R. Investigation

T. G. Investigation

T. R-F Investigation

A. P. Investigation

A. N Investigation and Formal analysis

M. B-S Project administration

A. F Conceptualization, Methodology, Validation, Investigation and Formal analysis

H. G Conceptualization, Methodology, Validation, Formal analysis, Resources, Supervision, and Funding acquisition

I. G Conceptualization, Methodology, Software, Validation, Formal analysis, Investigation, Resources, Writing, Visualization, Supervision, Project administration and Funding acquisition

## Supplementary Figures

**Figure S1:**
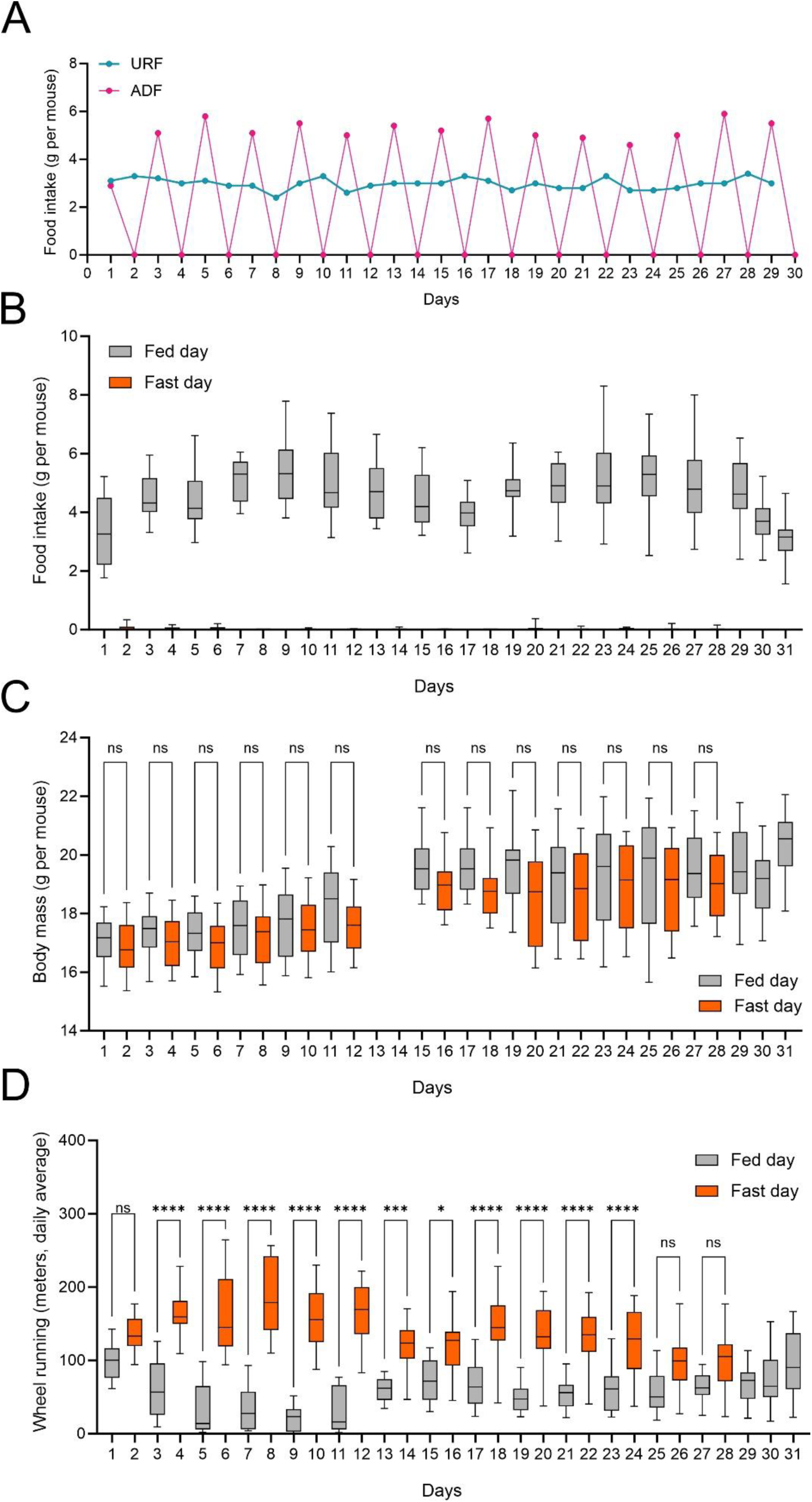
Metabolic parameters of ADF mice in fed and fasted days. **A.** Daily (24 h) food intake was measured, showing consistent intake in the URF group versus increased food intake in the ADF group during fed days (compensating for lack of intake in fasted days). **B.** Daily (24 h) food intake was measured in metabolic phenotyping cages, showing comparable results to conventional cages. **C.** Daily (24 h) body mass was measured in metabolic phenotyping cages, showing comparable results to conventional cages. **D.** Daily (24 h) wheel running was measured in metabolic phenotyping cages, showing increased wheel running in fasting days. - *P≤0.05, **P≤0.01, ***P≤0.001, ****P≤0.0001 by ordinary one-way ANOVA followed by Sidak post hoc analysis. Biological replicates: 16.

**Figure S2:**
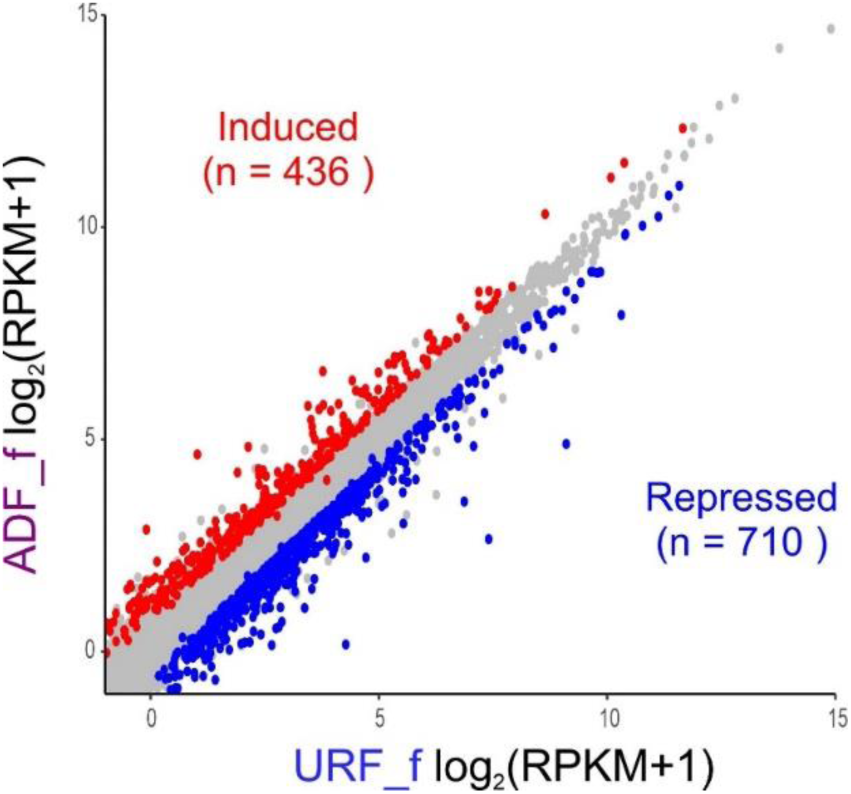
The transcriptional response to acute fasting differs significantly following ADF. Normalized mRNA expression values of genes regulated in ADF_f compared to URF_f. Red: induced genes; Blue: repressed genes; Grey: unchanged genes; Inclusion criteria for induction and repression: fold change ≥ 1.5, adj. p value ≤ 0.05; RPKM: reads per kilobase per million. - Data are presented as mean. Biological replicates: 3. Genes with RPKM < 0.5 were excluded.

**Figure S3:**
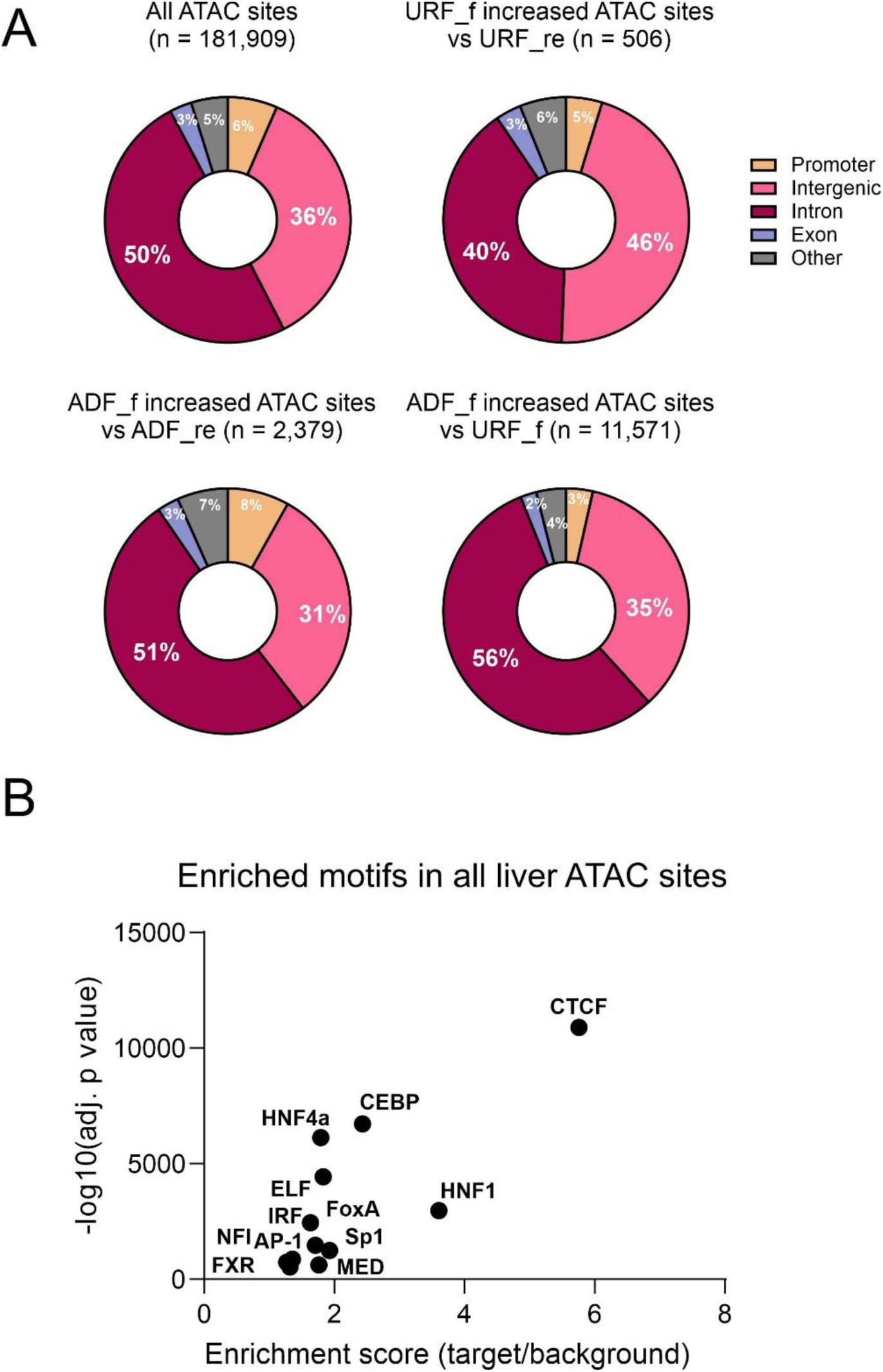
Most accessible sites harbor prototypical liver enhancer TF motifs and are not promoter-proximal. **A.** Genomic annotations of ATAC accessible sites show that the vast majority of liver accessible sites are not promoter-proximal. Promoter-proximal regions were defined as -1kb to +0.1kb from the transcription start site (TSS). **B.** Motif enrichment analysis of total ATAC accessible sites shows enrichment of liver lineage-determining factors known to bind hepatic enhancers, suggesting these sites are largely comprised of enhancers. - Biological replicates: 3. Motifs appearing in less than 1% of accessible sites were omitted (B).

**Figure S4:**
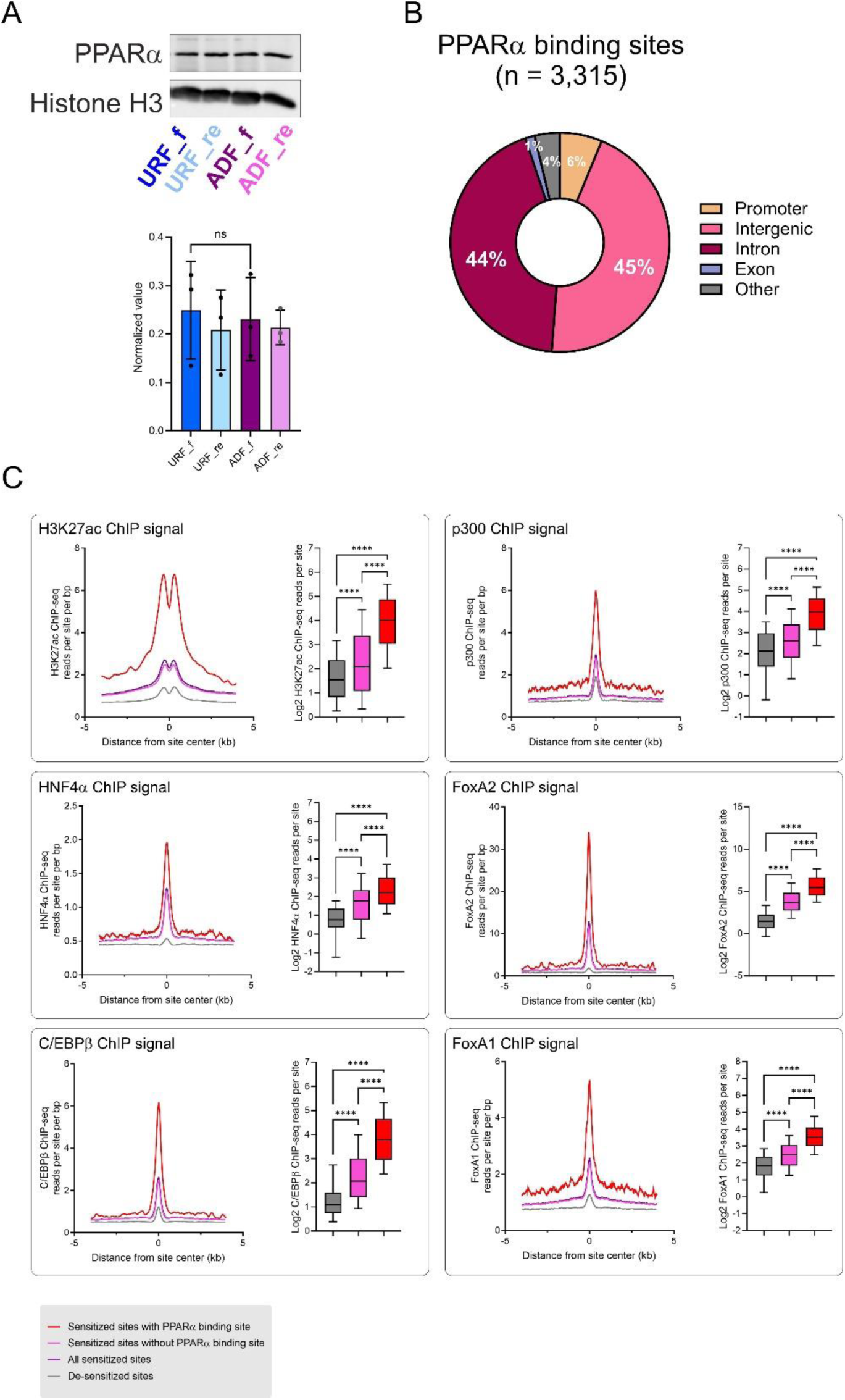
PPARα-bound sensitized sites are enriched with enhancer markers. **A.** The protein levels of PPARα were quantified using a western blot. **B.** Genomic annotations of PPARα binding sites show that the vast majority of sites are not promoter-proximal. Promoter-proximal regions were defined as -1kb to +0.1kb from the transcription start site (TSS). **C.** Quantification of ChIP-seq signal of various liver enhancer markers at sensitized and de-sensitized enhancers shows increased signal in sensitized enhancers which is further increased in PPARα-bound sensitized enhancers. ChIP-seq data was obtained from published datasets (see Methods). - Data are presented as mean ±S.D. *P≤0.05, **P≤0.01, ***P≤0.001, ****P≤0.0001 by ordinary one-way ANOVA followed by Sidak post hoc analysis by an unpaired two-tailed t-test. Biological replicates: 3 (A), 2 (B).

**Figure S5:**
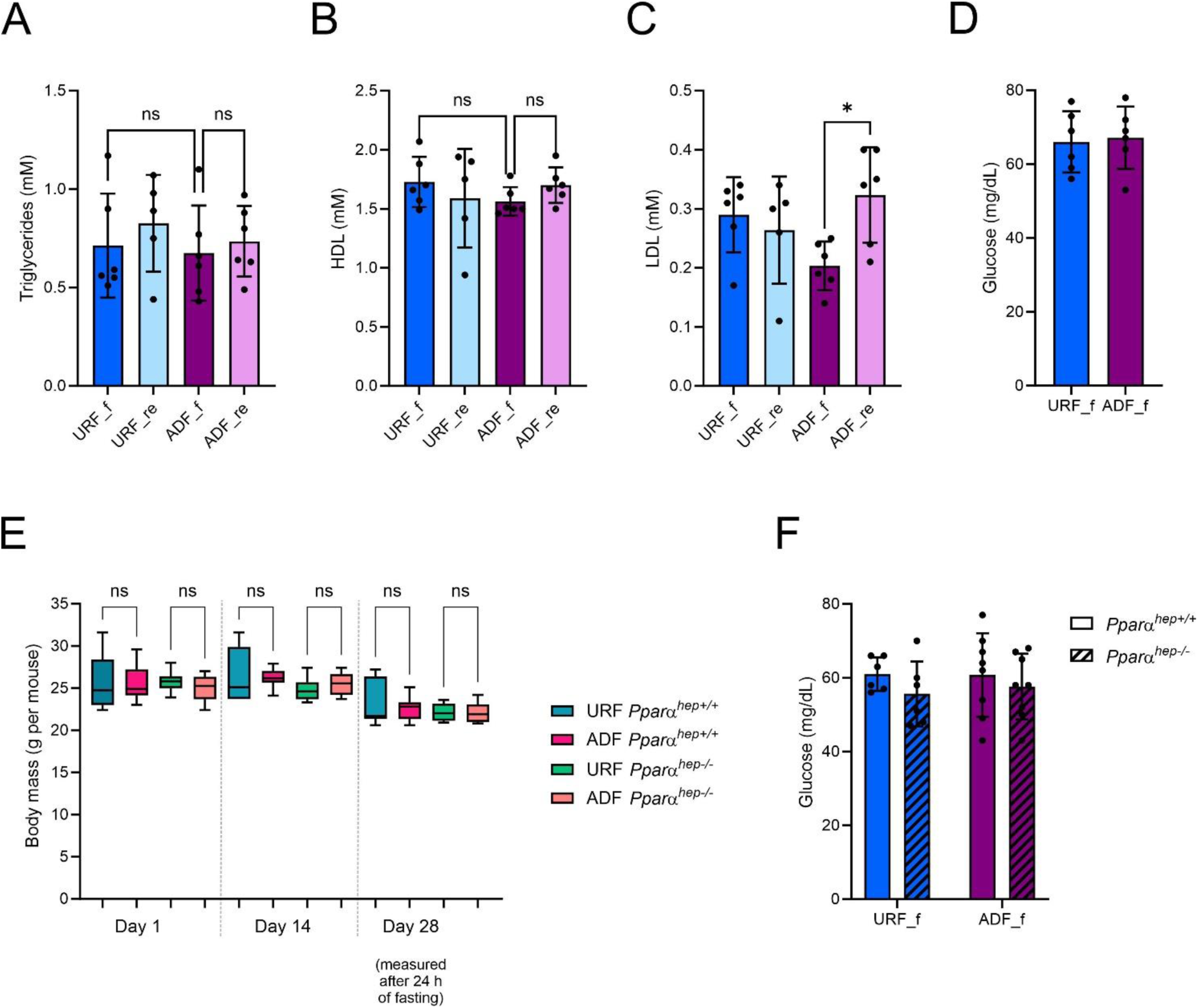
plasma parameters of ADF. **A-C**. The plasma levels of triglycerides, HDL and LDL were measured after 4 weeks of ADF, showing reduced levels of LDL following ADF. **D.** Blood glucose levels were measured following 24 h of fasting in ADF and URF mice, showing no effect of ADF on glycemia. **E.** Body mass was measured weekly in PPARα^hep+/+^ and PPARα^hep-/-^ mice, showing unchanged body mass in ADF mice as compared to URF mice. **F.** Blood glucose levels were measured following 24 h of fasting in PPARα^hep+/+^ and PPARα^hep-/-^ mice, showing no effect of ADF on glycemia. - Data are presented as mean ±S.D. *P≤0.05, **P≤0.01, ***P≤0.001, ****P≤0.0001 by ordinary one-way ANOVA followed by Sidak post hoc analysis by an unpaired two-tailed t-test. Biological replicates: 6-8 (A-C).

## Supplementary Table titles

<u>Sup. Table 1: Fasting-regulated genes in ADF and URF</u>

<u>Sup. Table 2: Sensitized and de-sensitized genes</u>

<u>Sup. Table 3: Fasting-regulated enhancers in ADF and URF</u>

<u>Sup. Table 4: PPARα binding sites and enriched motifs within them</u>

