## Supplementary figures and images for "Repeated fasting events sensitize enhancers, transcription factor activity and gene expression to support augmented ketogenesis"

### Figure S1

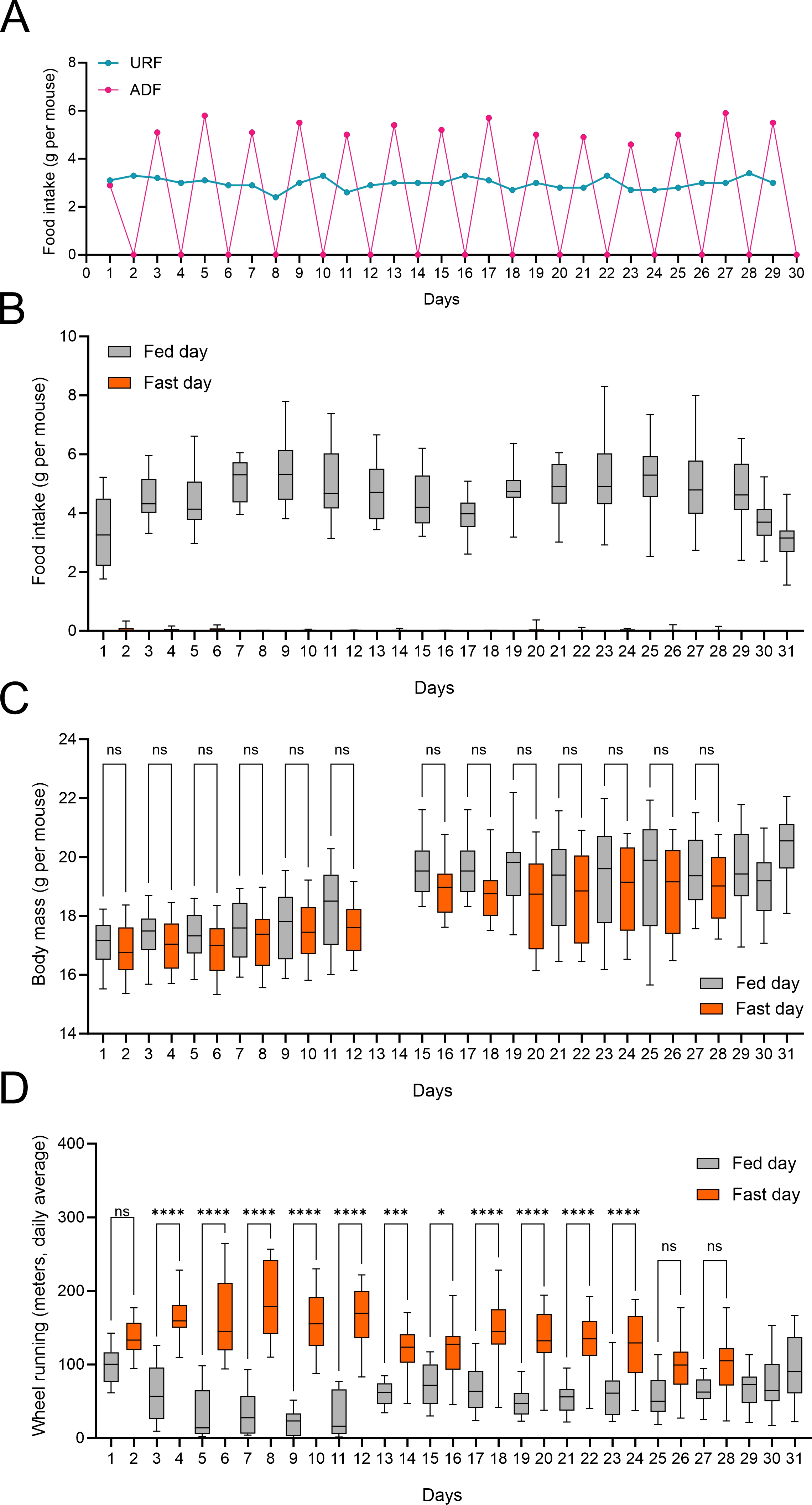

### Figure S2

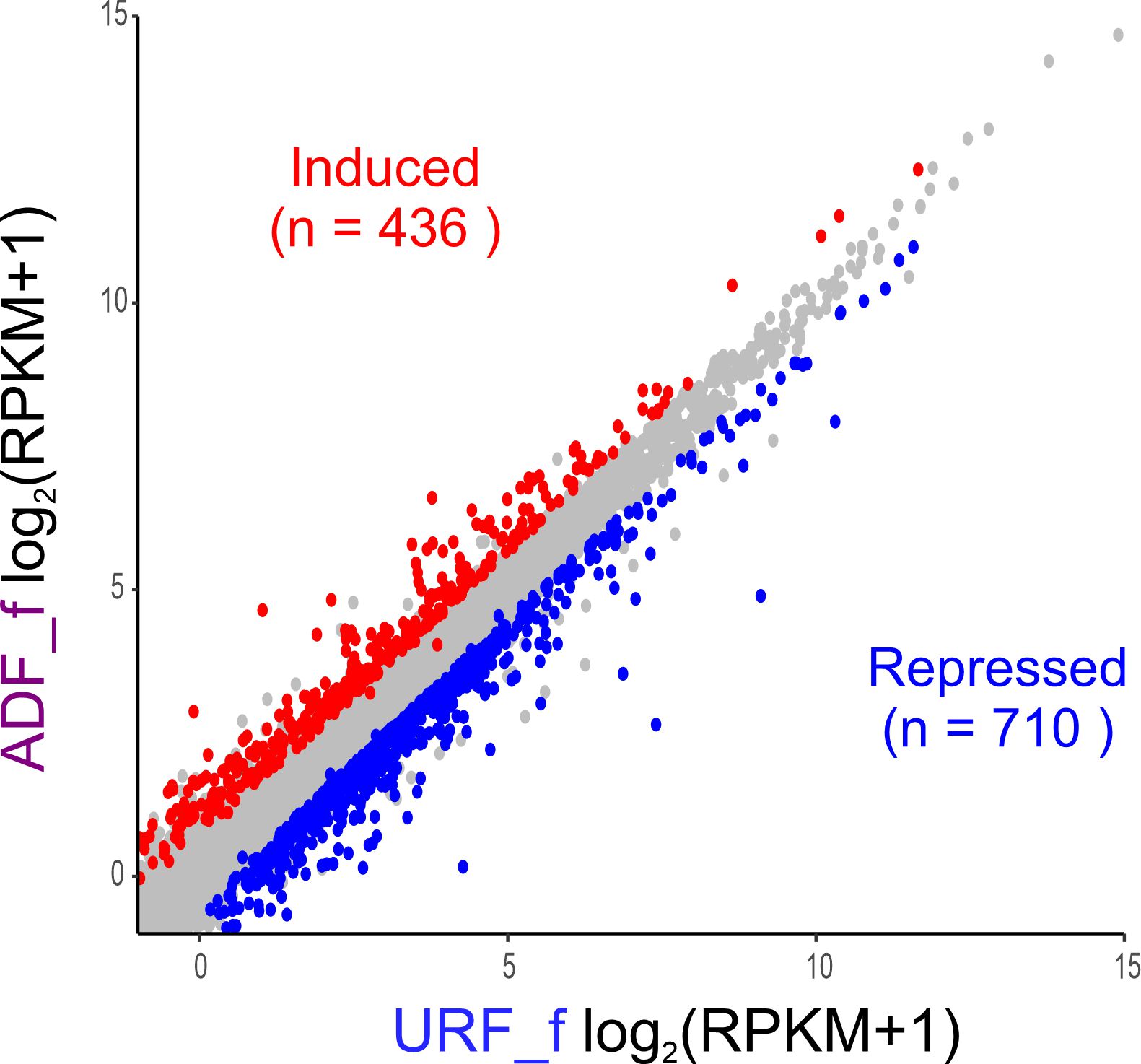

### Figure S3

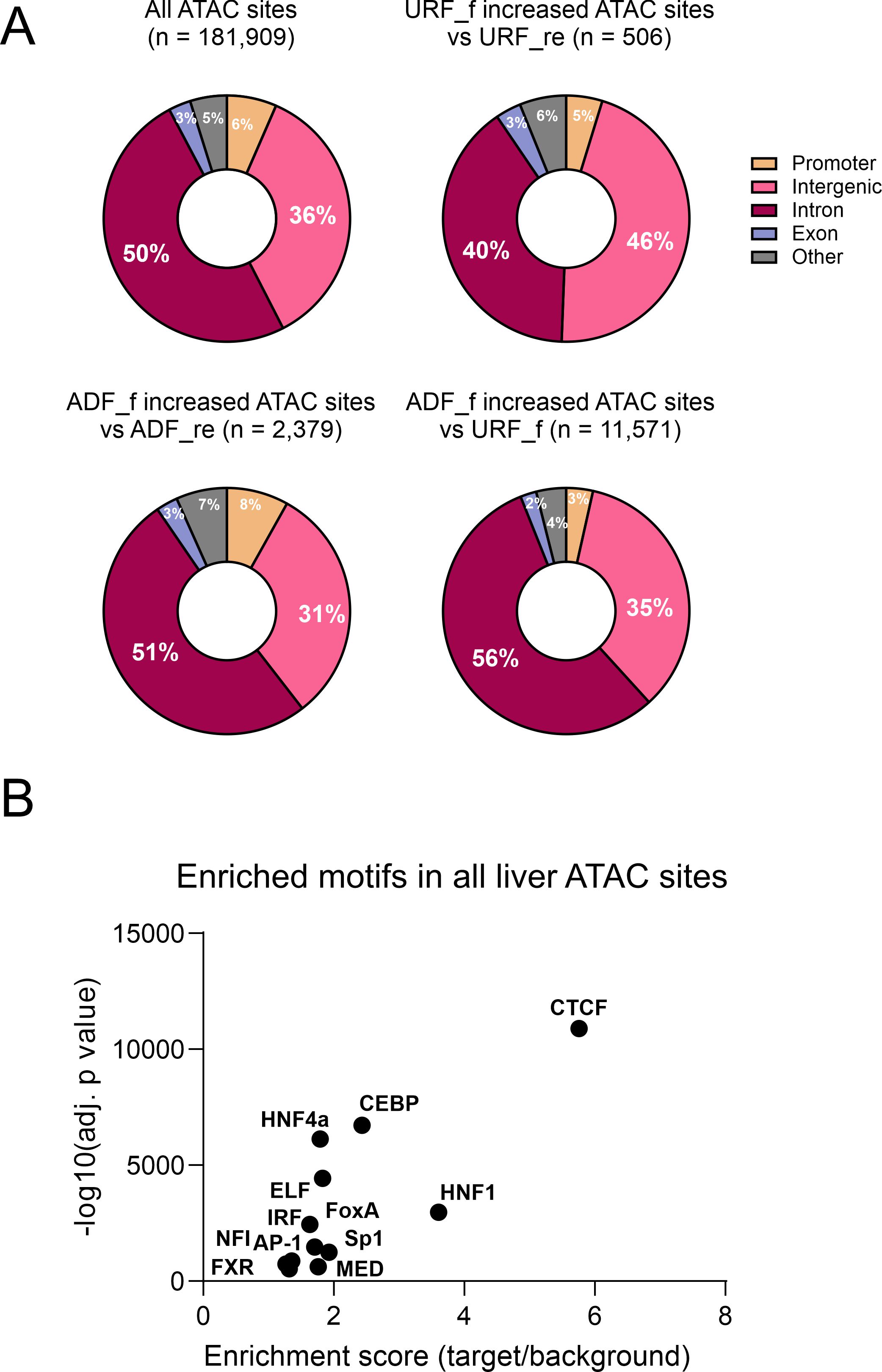

### Figure S4

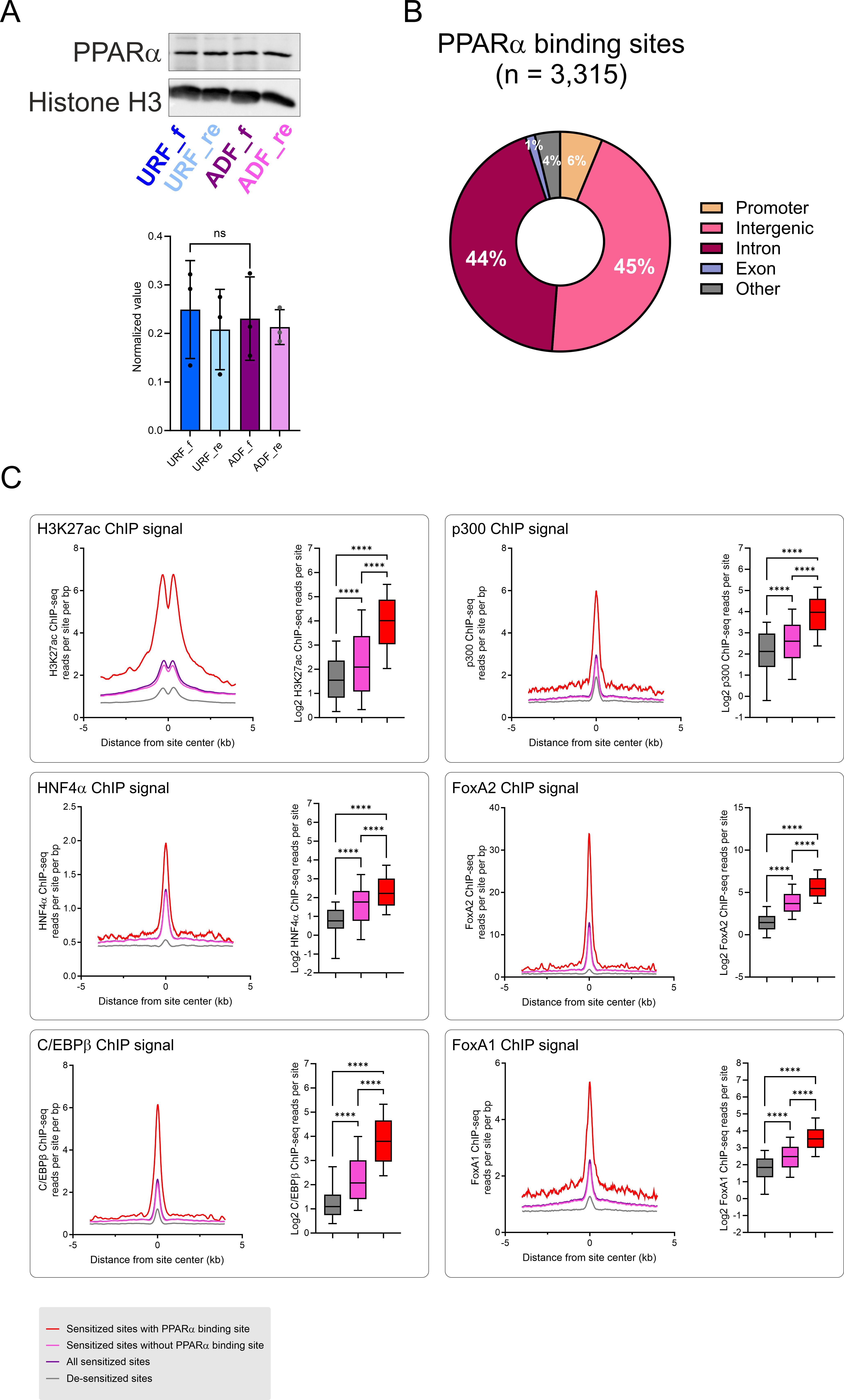

### Figure S5

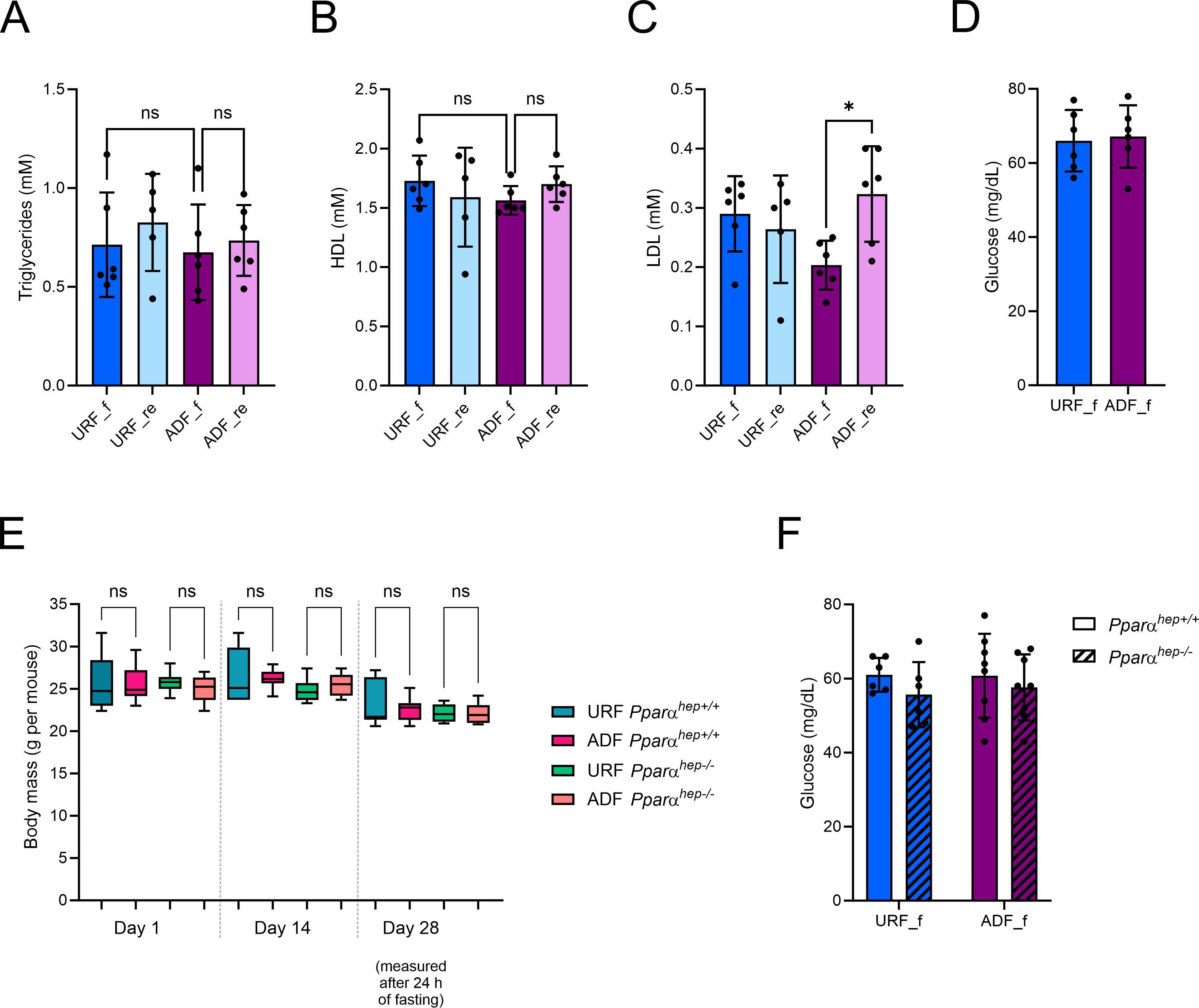
